# Single-cell profiling of environmental enteropathy reveals signatures of epithelial remodeling and immune activation in severe disease

**DOI:** 10.1101/2021.04.11.439202

**Authors:** Conner Kummerlowe, Thomas Wallach, Simutanyi Mwakamui, Travis K. Hughes, Nolawit Mulugeta, Victor Mudenda, Ellen Besa, Kanekwa Zyambo, Ira Fleming, Marko Vukovic, Ben A. Doran, Toby P. Aicher, Marc H. Wadsworth, Juliet Tongue Bramante, Amiko M. Uchida, Rabiah Fardoos, Osaretin E. Asowata, Nicholas Herbert, Henrik N. Kløverpris, John J. Garber, Jose Ordovas-Montanes, Zev Gartner, Alex K. Shalek, Paul Kelly

## Abstract

Environmental enteropathy (EE) is a subclinical condition of the small intestine that is highly prevalent in low- and middle-income countries. It is thought to be a key contributing factor to childhood malnutrition, growth-stunting, and diminished oral vaccine responses. While EE has been shown to be the by-product of recurrent enteric infection, its full pathophysiology remains unclear. Here, we mapped the cellular and molecular correlates of EE severity by performing high-throughput single-cell RNA-sequencing on 33 small intestinal biopsies from 11 adults with EE from Lusaka, Zambia (8 HIV-negative, 3 HIV-positive), 6 adults without EE in Boston, USA, and 2 adults from Durban, South Africa, which we complemented with published data from 3 additional South African adults from the same clinical site. By using these data to reanalyze previously-defined bulk-transcriptomic signatures of reduced villus height and decreased plasma LPS levels in EE, we found that these signatures may be driven by an increased abundance of surface mucosal cells – a gastric-like subset previously implicated in epithelial repair in the gastrointestinal tract. In addition, we identified several cell subsets whose fractional abundances associate with histologically determined EE severity, small intestinal region, and HIV infection. Furthermore, by comparing distal duodenal EE samples with those from three control cohorts, we identified dysregulated WNT and MAPK signaling in the EE epithelium and a T cell subset highly expressing a transcriptional signature of tissue-resident memory cells but with increased pro-inflammatory cytokine expression in the EE cohort. Altogether, our work illuminates epithelial and immune correlates of EE and provides new molecular targets for intervention.

**One Sentence Summary:** Using single-cell RNA-sequencing, we characterize the pathophysiology of environmental enteropathy (EE) – a highly prevalent condition of the small intestine that is thought to be a primary cause of global growth-stunting cases and a key contributing factor to childhood malnutrition and diminished oral vaccine responses – to derive insights into the epithelial and immune correlates of disease severity, suggesting new therapeutic targets for future investigation.

## INTRODUCTION

Environmental enteropathy (EE) is a subclinical condition of the small intestine that is driven by enteropathogen exposure through environmental contamination *(1, 2)*. Also referred to as Environmental Enteric Dysfunction (EED), EE impacts millions of children and adults around the world. It is associated with stunted growth, neurocognitive impairment, reduced oral vaccine efficacy, and life-long increased risk of metabolic syndrome *(1, 3, 4)*. Water, sanitation, and hygiene (WASH) interventions for preventing EE have proven ineffective, and ongoing work is assessing alternative therapeutic interventions such as antibiotics, anti-inflammatory therapeutics, and dietary supplementation *(5)*. However, development of effective treatments has been hindered by a limited understanding of the underlying mechanisms of EE.

Studies of the tissue biology of EE have been practically limited by operational constraints. Obtaining small intestinal biopsies from patients with EE requires esophago-gastroduodenoscopy (EGD) — a procedure that, while generally safe, carries increased risk to perform in undernourished pediatric populations. Accordingly, for ethical reasons, novel approaches and initial mechanistic studies using invasive techniques should be completed in affected adult populations before profiling pediatric patients. While the largest global health consequences of EE are seen in children, this condition also negatively impacts adult quality of life via its associated increased intestinal permeability, which can lead to systemic inflammation, increased risk of metabolic diseases, and reduced intestinal absorption *(1, 6)*. Given the environmental nature of enteric infections, finding unaffected controls for studies of EE in affected populations is similarly challenging. Thus, EE is often contextualized to health by either comparing intermediate EE with severe EE *(7)*, or by comparing EE patients to control cohorts in the United States or the United Kingdom *(8)*. The validity of these international comparisons is supported by the environmental nature of EE and the resolution of EE in Peace Corps volunteers upon repatriation to the United States *(9)*.

Pathologically, EE in the proximal small intestine is continuous, does not extend beyond the mucosa, and is characterized by reduced villus height, increased villus fusion, greater crypt depth, and increased microbial translocation *(10)*. However, in a study of Zambian children with EE and non-responsive growth stunting over time, reduced villus height was associated with decreased circulating levels of LPS (a measure of microbial translocation) *(11)*. A bulk transcriptomic study of Zambia children with enteropathy due to severe acute malnutrition (SAM) showed that many genes differentially upregulated in biopsies with reduced villus height were downregulated in biopsies from participants with high circulating levels of LPS *(7)*. These studies raise the intriguing possibility that EE is an adaptive response to potentially lethal enteropathogen exposure that comes at the cost of reduced absorptive capacity and thus impaired growth. However, a detailed picture of the cellular changes underpinning these processes is currently missing.

Histological analysis of EE has revealed increased abundance of lymphocytes (rather than granulocytes) and abnormalities of secretory cells, with reduced goblet cell numbers and abnormal Paneth cell morphology *(8, 12)*. Low plasma levels of tryptophan in children with growth stunting *(13)* and the amelioration of villus blunting in Zambian adults given tryptophan, glutamine, and leucine supplementation suggest that amino acid deficiency may play a role in epithelial remodeling in EE *(14)*. Bulk transcriptomic studies of EE duodenal biopsies, meanwhile, have revealed increased expression of NADPH oxidases, CXC chemokines, mucin genes, matrix metalloprotease genes, interferon stimulated genes, and antimicrobial genes including *LCN2, DUOX2*, and *DUOXA2 (15, 16)*. In addition, immunohistochemistry staining of the DUOX2 protein has been validated as a marker distinguishing tissues from Bangladeshi children with EE from control samples obtained from North American children with healthy tissue or with celiac disease *(16)*. Furthermore, transcriptomic analysis of feces from Malawian children has illustrated increased innate immune activity and interferon signaling, as well as reduced expression of mucins and pro-proliferative genes (including EGFR and MAPK7), in severe EE *(17)*. However, previous work has lacked the single-cell resolution required to localize these changes to specific epithelial and immune cell subsets in the small intestine.

High-throughput single-cell genomic profiling holds transformative promise for understanding the cellular populations, phenotypic states, and signaling changes that underlie EE *(18)*. In the small intestine, it is challenging to use bulk level measurements to resolve the states of rarer cell types (such as intestinal stem cells, enteroendocrine cells, or tissue resident lymphocytes) that play crucial roles in tissue maintenance *(19)*. In contrast, single-cell RNA-sequencing (scRNA-seq) can be used to comprehensively profile the distinct cellular subsets that compose complex tissues, thereby enabling analysis of rare yet influential cell subsets, and discovery of altered cell states and signaling. Illustratively, the unprecedented cellular resolution provided by scRNA-seq has been leveraged to identify novel features of human epithelial inflammation in the skin *(20)*, the nasal passages *(21)*, and the large intestine *(22)*. While scRNA-seq has also been used to study the healthy human small intestine *(23)*, to our knowledge, it has yet to be leveraged to understand EE.

Here, we applied the Seq-Well S^3^ platform for massively-parallel scRNA-seq *(24)* to profile small intestinal biopsies from 11 adults from a community in Zambia where EE is known to be ubiquitous *(25)*. Across these individuals, we profiled 27 biopsies spanning 3 small intestinal regions, HIV-positive and HIV-negative patients, and a range of histological EE severity scores. This provided capacity to define the cellular subsets associated with these covariates. In addition, by comparing EE biopsies with those from control groups from Durban, South Africa and Boston, USA, we identified upregulated of WNT signaling and downregulated of MAPK signaling in the EE epithelium and a T cell subset highly expressing a transcriptional signature of tissue-resident memory cells as well as pro-inflammatory cytokines in severe EE. In addition, relative to only the U.S. cohorts, the EE epithelium displayed reduced proliferative signaling and fractional abundances of goblet cells. Altogether, our data provide new insight into epithelial remodeling and immune signaling in EE, suggesting several novel therapeutic targets for further investigation.

## RESULTS

### scRNA-seq of the proximal small intestine with and without EE

We collected 27 small intestinal biopsies from 11 Zambian volunteers with EE (8 HIV-negative, 3 HIV-positive). For all 11, we profiled duodenal bulb and distal duodenum; for a subset, we also collected jejunal samples (**Table S1, S2**). Biopsies displayed varying levels of EE severity, with villus height:crypt depth ratios ranging from 0.64:1 to 2.38:1, compared to a normal ratio of 3:1 or greater (**Fig. 1A**, **Table S3**) *(8)*. Across these samples, we sought to identify the cellular correlates of intestinal region, HIV infection, and EE histological severity. However, due to both the widespread prevalence of EE in Zambia and a lack of existing screening methods to identify patients without EE, we could not obtain control biopsies from patients without EE in Zambia. Thus, we chose to distinguish the cellular and molecular features of EE biology by comparing to duodenal samples from 5 adults recruited from a gastrointestinal unit in Durban, South Africa (2 of which we profiled and 3 of which were studied previously using the Seq-Well S^3^ technology and the same tissue dissociation protocol *(26, 27)* as well as two cohorts of patients in Boston, USA where EE can safely be assumed not to occur: distal duodenum samples from 3 patients undergoing screening for eosinophilic esophagitis (EoE) and 3 patients undergoing duodenal resection for pancreatic malignancy (**Table S1, S2**). Among the former, pathology did not report eosinophilic gastroenteritis for any patients screened for EoE, and revealed that one patient did not have EoE; duodenal samples from pancreatic cancer patients meanwhile were taken from uninvolved resected tissue.

**Figure 1:**
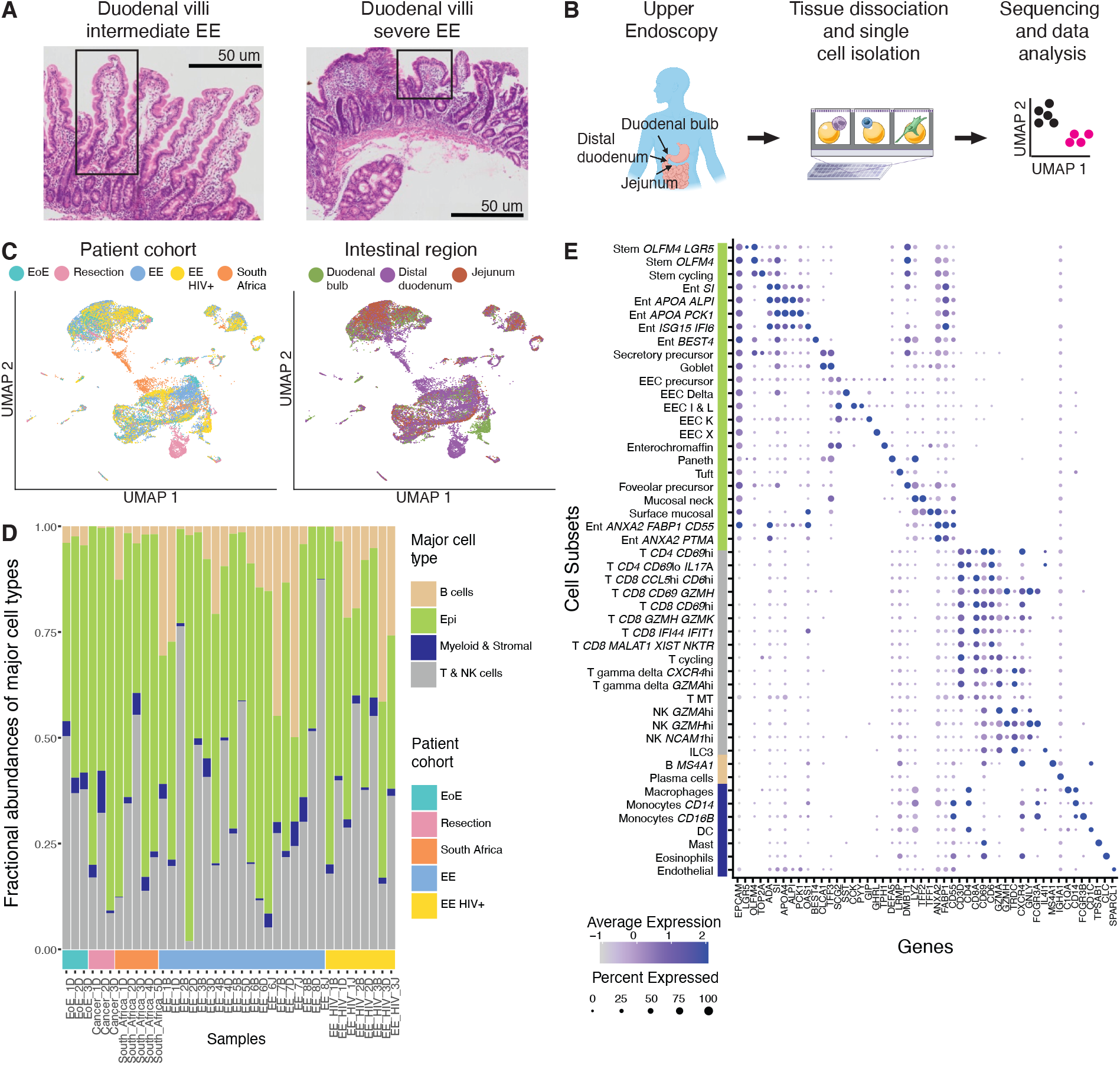
Single-cell RNA-sequencing of the upper small intestine with and without EE. **A,** H&E imaging of the small intestine with intermediate and severe EE. Intestinal villi with normal morphology (left image) and with severe blunting (right image) are highlighted in boxes. **B,** Experimental workflow: small intestinal biopsies from the duodenal bulb, distal duodenum, and jejunum were obtained via endoscopy or tissue resection, dissociated into cells, loaded onto a Seq-Well array, processed for single-cell sequencing, and analyzed. **C,** UMAP visualization by patient cohort and intestinal region for all 26,556 high quality cells from 38 samples and 22 patients. **D,** Fractional abundances of major cell types amongst all single cells analyzed per sample. **E**, Expression of marker genes for final cell subsets. Dot size represents the fraction of a cell subset (rows) expressing a given gene (columns). Dot hue represents the scaled average expression by gene column. For clarity, dots for genes expressed in 5% or less of cells within a given subset are not shown.

### Iterative clustering identifies cell subsets across heterogenous patient conditions

In total, we analyzed 26,556 high quality single-cell transcriptomes across 38 samples from the Zambian, U.S., and South African cohorts (**Fig. 1B**). After data pre-processing (see **supplementary methods**), UMAP visualization of the entire dataset revealed differences in cellular distribution by patient cohort, intestinal region, and HIV status (**Fig. 1c**). In order to rigorously identify cellular subsets across these biological covariates, we applied an existing pipeline for iterative clustering of cell subsets to the Zambian and U.S. datasets (which were available before the South African data and processed in a different laboratory) *(28, 29)* annotating the resulting clusters by known lineage markers and, in rare cases, merging together subsets across batches that co-expressed known lineage markers (see **supplementary methods**). Next, we integrated this data with the South African data, using the Seurat V3 method to correct for potential batch effects between data collected in different laboratories while allowing for the possibility that new cell subsets could exist in the South African data *(30)*. These analyses revealed that the expected major cell types (epithelial cells, T & NK cells, B cells, myeloid & stromal cells) were represented in almost all biopsies and that epithelial cells were the most abundant major cell type (**Fig. 1D**). Along with more standard QC metrics (**Fig. S1-S4**), this indicated consistent sample quality. We then identified 48 detailed cellular subsets that varied in abundance across samples (**Fig. 1E, Fig. S1-S4, Table S4**) *(19, 22, 23)*. These 48 subsets included: 23 epithelial subsets – inclusive of 3 stem cell subsets, 1 subset highly expressing *BEST4*, 1 secretory precursor subset, 1 goblet cell subset, 1 Paneth cell subset, 1 tuft cell subset, 6 enteroendocrine subsets, 1 surface mucosal cell subset (high expression of *TFF1* and *MUC5AC)*, 1 mucosal neck cell subset (high expression of *TFF2* and *MUC6)*, one foveolar precursor subset, and 6 enterocyte subsets, two of which highly expressed *ANXA2* (a marker of intestinal epithelial cells engaged wound healing *(31)*) and either *FABP1* or *PTMA* (markers of dedifferentiating intestinal cells in a murine model system *(32)*); 16 T & NK cellular subsets – consisting of 5 *CD8*^hi^ T cell, 2 *CD4^hi^* T cell, 2 γδ T cell, 3 NK cell, 1 ILC3, and 3 T cell subsets respectively defined by interferon genes, mitochondrial genes, and cell cycle genes; 6 myeloid subsets – inclusive of macrophages, *CD14* monocytes, *CD16* monocytes, dendritic cells, mast cells, and eosinophils; 2 B cell subsets – plasma cells and *MS4A1* (i.e., CD20) expressing B cells; and, one stromal cell type, endothelial cells.

### Comparison to past bulk RNA-seq EE signatures implicates surface mucosal cells expressing *DUOX2* in epithelial remodeling in EE

When identifying cell subsets, we noticed a striking similarity between the marker genes of surface mucosal cells (a cell subset most commonly found in the distal stomach that highly expresses *TFF1* and *MUC5AC* and has been implicated in the mucosal injury response across the gastrointestinal tract) *(33)* and existing bulk transcriptomic gene signatures characterizing EE samples with reduced villus height and decreased plasma LPS levels in Zambian children with SAM (**Fig. S5A**) *(7)*. Scoring all subsets on these signatures, we found that surface mucosal cells were significantly enriched for both signatures relative to all other cell subsets (p < 10^15^, Wilcoxon test) (**Fig. 2A, B, Fig. S5B, C**). This suggests that a higher abundance of this cell subset may have driven these previously derived signatures. To examine this possibility more closely, we visualized the overlap between the genes from these signatures and the marker genes of all subsets (**Fig. 2C, D**). This revealed that the inferred enrichment of surface mucosal cells in this these signatures is driven by high expression of known surface mucosal cell markers *(34)*, as well as the antimicrobial genes *LCN2, DUOXA2*, and *DUOX2*. These last three genes were also highly upregulated in EE relative to controls in a recent bulk transcriptomic study of Bangladeshi children *(16)*. That study demonstrated that immunohistochemical staining of DUOX2 protein in the villus epithelium distinguished EE biopsies from controls. Given that reduced villus height in EE has been shown to associate with decreased microbial translocation over time *(11)*, it is remarkable that surface mucosal cells scored highly for signatures of both reduced villus height and decreased microbial translocation (as measured by decreased plasma LPS concentrations). This suggests that this cell subset may be associated with adaptive shortening of the villus in response to enteropathogen exposure.

**Figure 2:**
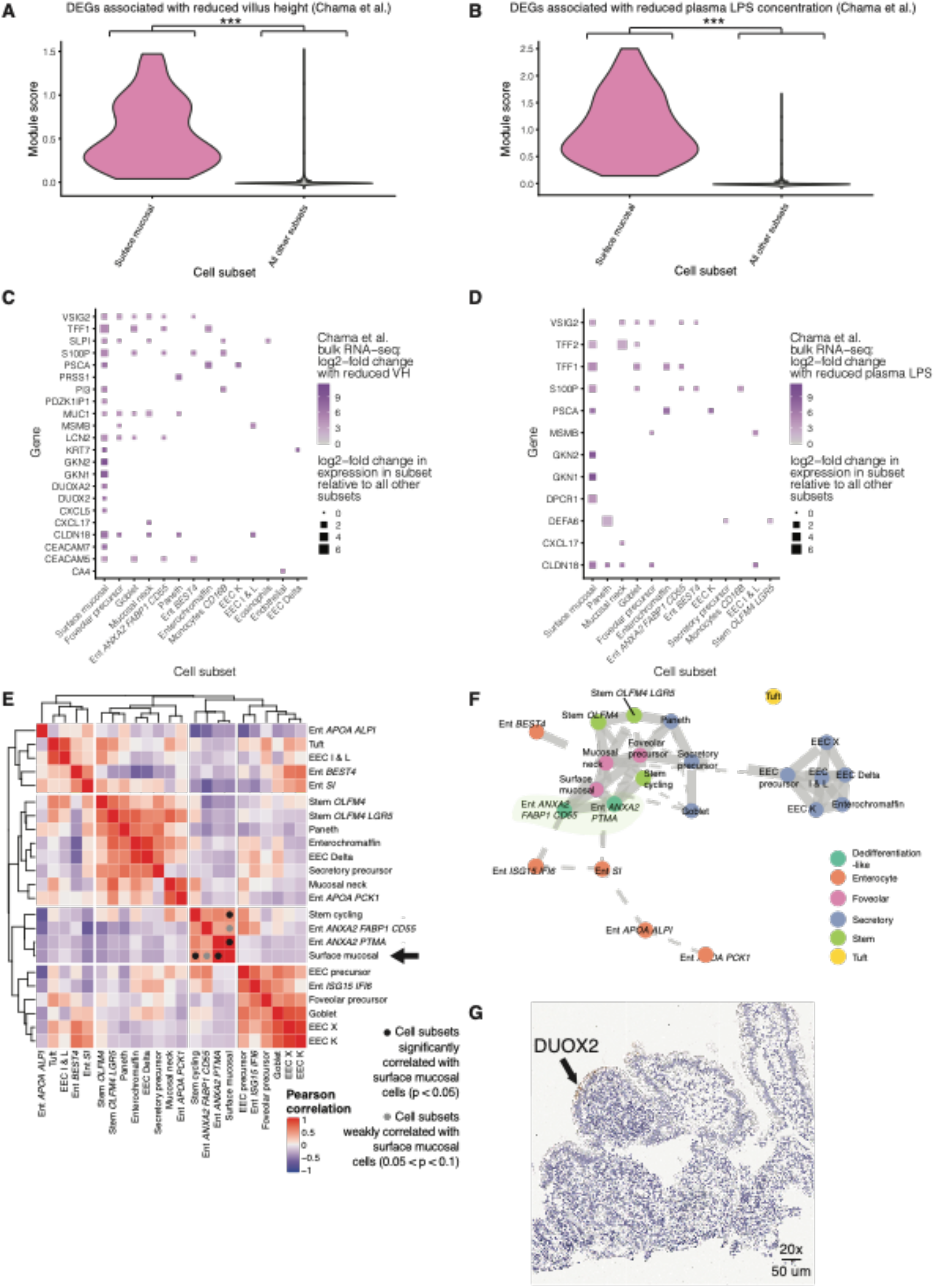
Comparison to past bulk RNA-sequencing EE signatures implicates surface mucosal cells expressing DUOX2 in epithelial remodeling in EE. **A,** Violin plot of a module score generated from genes differentially upregulated in bulk RNA-sequencing of samples with EE and reduced villus height (VH) in Chama et al. *(7)* Surface mucosal cells were enriched for this signature relative to all other subsets (***, p < 0.001; Wilcoxon test) **B,** Violin plot of a module score generated from genes differentially upregulated in bulk RNA-sequencing of samples with EE and decreased plasma LPS concentrations in Chama et al. *(7)* Surface mucosal cells were enriched for this signature relative to all other subsets (***, p < 0.001; Wilcoxon test) **C,** Dot plot of cell subset marker genes that overlapped with the genes used to generate the module scores in panel a **D,** Dot plot of cell subset marker genes that overlapped with the genes used to generate the module scores in panel b. **E,** Hierarchically clustered heatmap of the Pearson correlations between the fractional abundances of epithelial cells within duodenal bulb samples from patients with EE. Cell subsets significantly correlated with surface mucosal cells are highlighted with a black circle (Permutation testing, p < 0.05). Cell subsets weakly correlated with surface mucosal cells are highlighted with a grey circle (0.05 < p < 0.1; Permutation testing). **F,** Partition-based graph abstraction (PAGA) trajectory visualization of epithelial subsets. **G,** H&E (purple) and immunohistochemical staining for DUOX2 protein (brown) on an EE biopsy from the duodenal bulb.

Next, we sought to identify the epithelial subsets whose fractional abundances correlated with those of surface mucosal cells in our data. While the *DUOX2* signature was present in distal duodenal samples in past bulk transcriptomic studies, in our data, surface mucosal cells expressing *DUOX2* predominantly came from duodenal bulb samples of patients with EE (**Fig. S2C**). Thus, we focused our analysis on the duodenal bulb. We correlated the fractional abundances of all subsets in these samples and hierarchically clustered the results (**Fig. 2E**). Surface mucosal cell fractional abundances were significantly correlated (Permutation testing, p < 0.05) with those of cycling stem cells (Stem cycling) and the Ent *ANXA2 PTMA* subset which highly expressed marker genes of dedifferentiating enterocytes (*PTMA*) *(32)* and wound associated epithelial cells (*ANXA2*) *(31)* in murine studies. In addition, surface mucosal cell fractional abundances were weakly correlated with those of the Ent *ANXA2 FABP1 CD55* subset (which also highly expressed marker genes of dedifferentiation and wound healing) (Permutation testing, p < 0.1). This suggests that surface mucosal cells are associated with a wound healing-like epithelial state.

To further characterize the Ent *ANXA2 PTMA* and Ent *ANXA2 FABP1 CD55* subsets that cooccurred with surface mucosal cells, we inferred differentiation trajectories for the epithelial cells in our dataset with PAGA *(35)* – an unbiased algorithm for estimating cell trajectories based on cell subset similarity (**Fig. 2F**). These two epithelial subsets and surface mucosal cells all lay in between mature enterocyte and stem cell subsets in the inferred differentiation hierarchy. In agreement, upon running the RNA velocity package Velocyto, these subsets again lay in between the enterocyte and stem cell subsets (**Fig. S5D, E**) *(36)*. Together, these results suggest that the presence of these wound healing-like epithelial subsets and *DUOX2* expressing surface mucosal cells in EE is associated with a shift towards an intermediate-like epithelial phenotype. This is consistent with the fact that surface mucosal cells highly express an array of transcripts encoding mucins and antimicrobial proteins, including *DPCR1* (also known as *MUCL3*), which has been shown to be a transcriptomic marker of wound associated epithelial cells in murine studies (**Table S4**) *(37)*. To visually contextualize surface mucosal cells in the epithelium, we immunohistochemically stained samples for DUOX2 protein and found that DUOX2 localized to the villus tip in blunted villi (**Fig. 2G, Fig. S6**). As only surface mucosal cells express *DUOX2*, this further suggests that surface mucosal cells are associated with an intermediate wound healing-like phenotype that occurs in response to enteropathogen mediated damage. This possibility would provide a mechanism to explain the previous observation *(11)* that, over time, villus shortening is associated with decreased microbial translocation in EE.

Furthermore, surface mucosal cells uniquely expressed *MUC5AC*, whose protein product has been shown to colocalize with *H. pylori* in gastric infections (**Fig. S5A, Table S4**) *(38)*. Thus, we hypothesized that the presence of surface mucosal cells in the EE cohort may have been associated with ongoing *H. pylori* infection. To address this possibility, we employed the metagenomic classification tool Kraken 2 to quantify the number of reads mapping to the *H. pylori* transcriptome in the sequencing data *(39)*. Relative to samples from the control cohorts from the U.S. and South Africa, 6 samples from 4 participants in the EE cohort contained significantly higher levels of *H. pylori* transcriptome mapping reads, suggesting that a *H. pylori* infection was ongoing in these participants (**Fig. S5F**). In addition, relative to samples from the distal duodenum and jejunum, two samples from the duodenal bulb contained significantly higher numbers of *H. pylori* transcriptome mapping reads (**Fig. S5G**). Thus, our data suggest that the presence of surface mucosal cells and the associated epithelial intermediate wound healing-like phenotype in only samples from the duodenal bulb from participants in the EE cohort may be associated, in part, with *H. pylori* infection in these samples.

### Mapping the cellular correlates of intestinal region and disease severity in HIV-negative EE

Next, we sought to leverage the single-cell resolution of our scRNA-seq dataset to examine previously undescribed yet fundamental variation in cellular subsets associated with small intestinal region and histologically determined EE severity. To avoid any confounding impacts of HIV infection biology, we focused this analysis on the eight HIV-negative participants with EE in our study.

First, we identified cell subsets whose fractional abundances robustly shifted across intestinal region within HIV-negative EE patients (see **Supplementary Methods, Fig. 3A**). Duodenal bulb samples were enriched for surface mucosal cells, mucosal neck cells, and intermediate wound healing-like enterocytes highly expressing *ANXA2, FABP1*, and *CD55*, as well as three T cell subsets expressing markers of immune activation *(IL17A, CXCR4, GZMA) (40, 41)*. On the other hand, distal duodenal samples were enriched for enterocytes, goblet cells, and stem cells highly expressing *OLFM4*. Due to a limited number of jejunal samples in our study, no cell subsets were significantly and robustly enriched in jejunal samples. These changes reveal that the duodenal bulb contains a more gastric-like transcriptional program than the distal duodenum in EE. This suggests variation in the cellular features of EE along the proximal small intestine.

**Figure 3:**
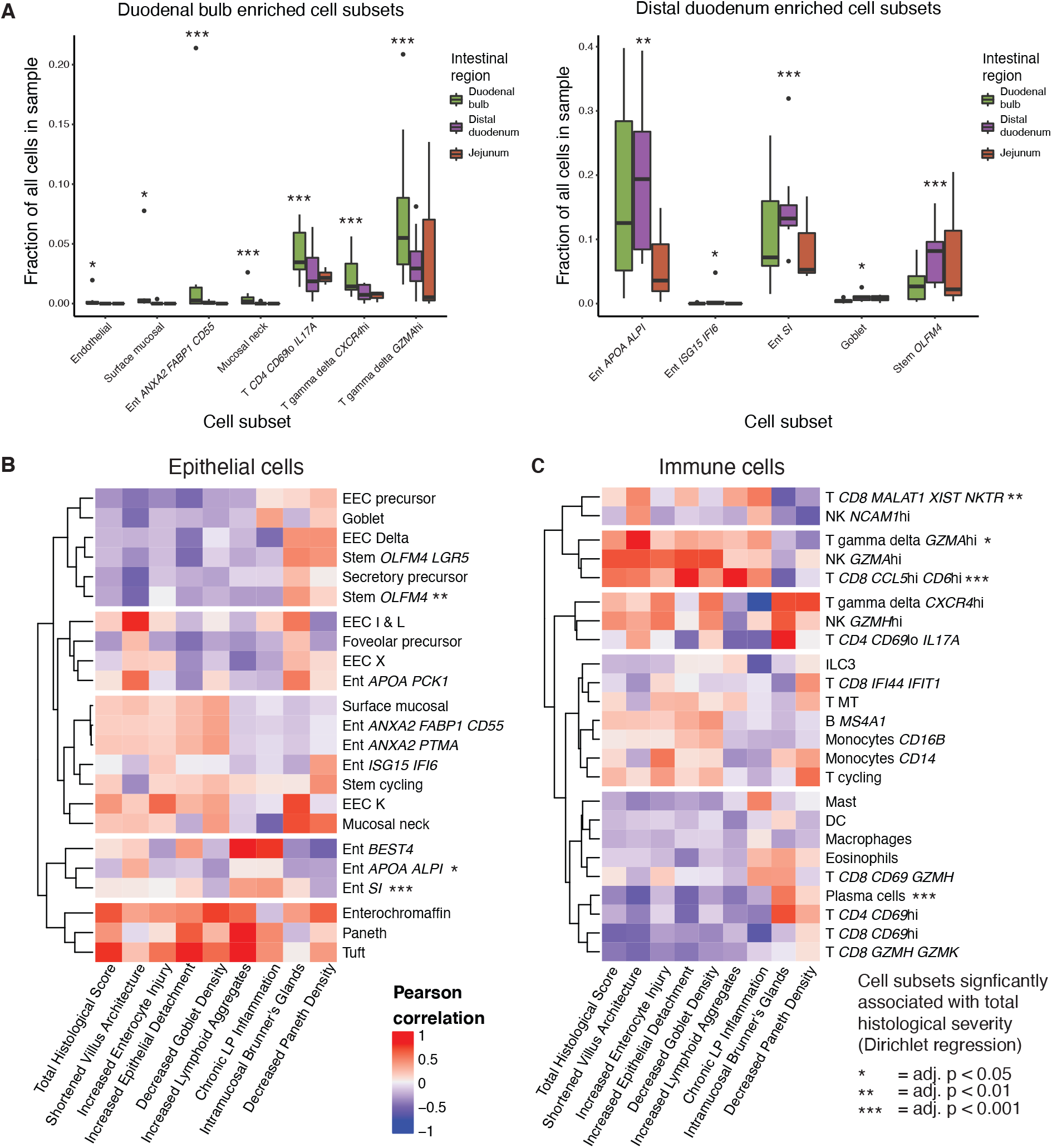
Identifying cell subsets associated with intestinal region and histologically determined EE severity within HIV-negative EE patients. **A,** Cell subsets enriched in duodenal bulb and distal duodenal samples from HIV-negative EE patients (*, adj. < 0.05; **, adj. p < 0.01; ***, adj. p < 0.001). No subsets were enriched in jejunal samples from HIV-negative EE patients. **B,** Hierarchically clustered heatmap of HIV-negative EE epithelial cell subset relative abundance Pearson correlations with component scores of the total EE histological severity score. In order to account for compositional dependencies in the data, significant relationships were determined with Dirichlet regression (*, adj. p < 0.05; **, adj. p < 0.01,; ***, adj. p < 0.001). **C,** Hierarchically clustered heatmap of HIV-negative EE immune cell subset relative abundance Pearson correlations with component scores of the total EE histological severity score. Significant relationships determined with Dirichlet regression (*, adj. p < 0.05; **, adj. p < 0.01; ***, adj. p value < 0.001).

Second, we sought to identify correlates of histologically determined EE severity across the entire small intestine. For eleven biopsies with matched histology and severity scores (**Table S5**) *(8)*, we used a Dirichlet regression model to assess how severity associated with cell subset composition and regress out small intestinal region-specific effects (**Fig. 3B,C**). As Dirichlet regression accounts for compositional dependencies in sequencing data, significant subsets do not always display the greatest absolute Pearson correlations with total EE severity, as these were generally observed in rarer subsets (e.g., tuft cells) whose relative abundances were likely to be highly influenced by random sampling or by shifts in the abundances of other subsets. Across small intestinal regions, in the epithelium, greater severity was significantly associated with lower fractional abundances of mature enterocytes and stem cells highly expressing *OLFM4*, as well as higher fractional abundances of immature enterocytes. This indicates that more severe EE may lead to a lower abundance of progenitor and mature cell subsets and a higher abundance of intermediate-like cell subsets. In the immune compartment, greater severity associated with a higher abundance of two T cell subsets expressing markers associated with inflammation in the intestine (*GZMA* and *CD6*) *(40)* and one T cell subset with high expression of *MALAT1* and the lowest median number of UMIs of all T cell subsets, which together suggest that this subset may represent low quality pre-apoptotic cells (**Fig. S2D**) *(42)*. In line with past histological findings in children with EE in the Gambia *(43)*, plasma cells abundances decreased with EE severity. Altogether, these results suggest that severe EE is characterized by an intermediate-like epithelial phenotype and provides new resolution into the lymphocyte subsets associated with EE.

### Identifying the cellular features associated with both HIV infection and EE

Next, we sought to characterize the impact of highly active antiretroviral treated HIV infection on EE pathology. First, we examined the subset of the EE dataset (16 biopsies from 8 patients including all 3 HIV-positive patients) for which we obtained matched histological images and found that HIV-positive samples displayed higher EE severity than HIV-negative samples (Wilcoxon test, p = 0.034) (**Fig. S7A**). Next, we examined robust shifts in cell subset fractional abundances with HIV infection (**Fig. S7B, C**). Significant changes included known features of HIV biology, such as decreased fractional abundances of *CD4*^hi^ T cells *(27, 44)* and increased fractional abundances of γδ T cells highly expressing the HIV co-receptor *CXCR4 (45)*. In addition, within duodenal bulb samples, HIV pathology associated with increased fractional abundances of intermediate wound healing-like enterocytes highly expressing *ANXA2, FABP1*, and *CD55* (**Fig. S7D**). This suggested that HIV pathology may also contribute to the presence of these subsets within the duodenal bulb. Altogether, these results suggest that HIV infection has a distinct influence on EE pathology.

### Inflammation in EE is accompanied by an immature epithelial phenotype and pro-inflammatory tuft cell gene expression changes

We next sought to identify features that distinguished EE small intestine samples from those without EE. To accomplish this, we compared the HIV-negative EE distal duodenal samples with matched small intestinal samples from participants in Durban, South Africa and two groups of matched small intestinal samples from Boston, USA: distal duodenal samples from patients referred to endoscopy for suspected EoE and distal duodenal samples from surgically resected uninvolved tissue from patients with pancreatic cancer. While histology was not available for the pre-existing South African dataset, H&E staining of a duodenal biopsy from a separate patient at the clinical site in Durban revealed features of EE, including villus blunting, goblet cell depletion, and Paneth cell depletion, but no signs of inflammation or lymphocyte infiltration (**Fig. S8A**). This raised the concern that the scRNA-seq samples from this site may also display features of EE. To investigate this possibility, we performed a pairwise comparison of the fractional abundances of all cell subsets between the three geographical locations in this study: Zambia, South Africa, and the U.S (**Fig. S8B-D**). Relative to the U.S. cohorts, both the Zambian and South African cohorts have reduced goblet cell and increased plasma cell fractional abundances. As previous findings have shown that these populations are reduced in EE relative to healthy controls, this suggested that both the Zambian and South African cohorts may display some features of EE *(8, 46)*. However, plasma cells and T cell subsets expressing markers of inflammation *(IL17A, GZMA)* were increased in fractional abundance in the Zambia EE cohort relative to the South African cohort *(40, 47)*. This suggested that not all features of EE are apparent in the South African samples. Thus, we took two approaches towards identifying distinguishing attributes of confirmed EE in the Zambian cohort. First, we took a conservative approach that minimized the risk of cohort or geographic specific effects dominating our analysis by comparing the Zambian cohort to all other cohorts from the U.S. and South Africa. Then, we compared the Zambian cohort with confirmed EE to only the U.S. cohorts in order to identify any differences that may have been masked in the previous analysis due to any potential features of EE in the South African cohort. In all these comparisons, we focused on biological features that: 1) were differentially regulated in EE compared to all outgroups and 2) displayed the same direction of regulation when comparing EE samples to each outgroup individually.

Comparing epithelial cells from patients with confirmed EE with those from both the U.S. and South Africa, we identified shifts in cell subset proportions, including an enrichment of stem cells highly expressing *OLFM4*, foveolar precursor cells, and enterocytes co-expressing *APOA4* and *ALPI* in EE relative to controls; and reduced fractional abundance of EEC K cells relative to controls (**Fig. 4A**). Overall, this suggested that the EE epithelium may be biased towards immature cellular subsets. Accompanying these changes, epithelial compartment-wide and cell-type specific differential expression between EE and control cohorts revealed upregulation of genes (*PIGR, DMBT1, CCL25, REG1A, SLC39A14*) involved in mucosal defense mechanisms (antibody transport, NOD2 signaling, lymphocyte recruitment, and ion transport, respectively) (**Fig. 4B,C Table S6**) *(48–51)*. Goblet cells and secretory precursor cells both highly upregulated *REG3A* in EE relative to controls (**Fig. 4C**), which is consist with an immature goblet cell phenotype *(52)*. EE samples also displayed compartment-wide upregulation of *CTNNB1*, a key component of WNT/β-catenin signaling. In agreement, inferring pathway signaling activity in EE with PROGENy revealed increased WNT signaling in all three stem cell subsets and decreased MAPK signaling in cycling stem cells and stem cells highly expressing *OLFM4* (**Fig. 4D**) *(53)*. As WNT signaling has been shown to suppress MAPK induced proliferation and differentiation of intestinal stem cells into mature cell subsets, our results suggest that mucosal defense in EE is accompanied by epithelial signaling promoting an immature phenotype *(54)*.

**Figure 4:**
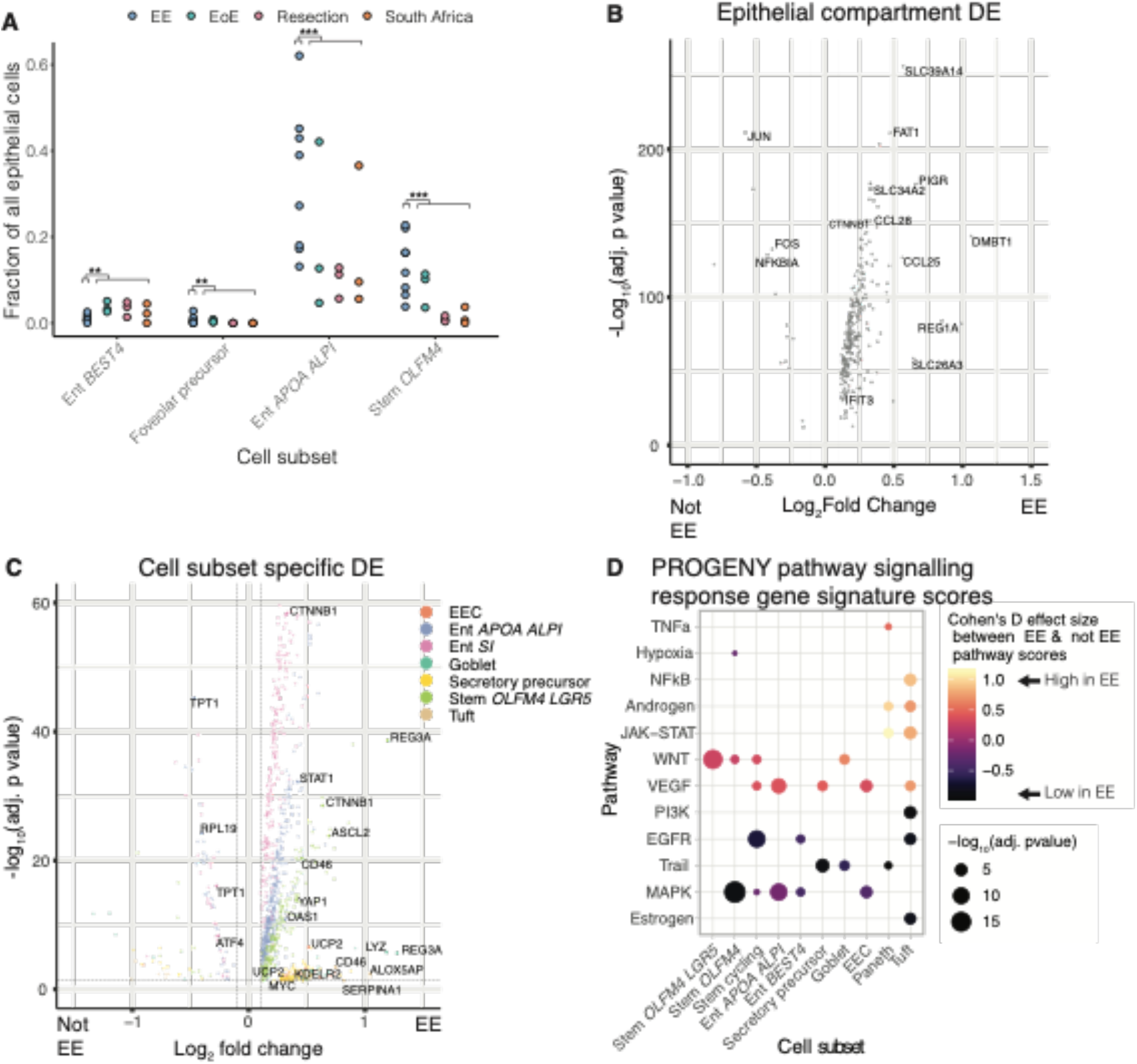
The epithelium of EE is characterized by increased interferon response, increased WNT signaling, and decreased MAPK signaling. **A,** Cell subsets with significant shifts in relative abundances between EE and all three control cohorts. (*, adj. p < 0.05; **, adj. p < 0.01; ***, adj. p < 0.001; Fisher’s exact test). **B,** Genes differentially expressed in the epithelial compartment in EE relative to all three control cohorts. Horizontal and vertical dashed lines respectively refer to an adjusted p value threshold of 0.01 and a log fold change threshold of 0.1. **C,** Genes differentially expressed in EE relative to all three control cohorts within specific cellular subsets. Horizontal and vertical dashed lines respectively refer to an adjusted p value threshold of 0.01 and a log fold change threshold of 0.1. **D,** PROGENy pathway prediction scores for epithelial cells in EE relative to controls

In agreement, stem cells highly expressing *OLFM4* and *LGR5* highly expressed *YAP1* – a portion of the Hippo pathway that negatively regulates stem cell differentiation into mature cells *(55)*. The upregulated activity of these signaling pathways may be the cause or consequence of the villous blunting that is a hallmark of EE pathogenesis. In addition, tuft cells in EE highly expressed *ALOX5AP* (which plays a role in inflammation in asthma, arthritis, and psoriasis via leukotriene biosynthesis *(56)*) and *SERPINA1* (which encodes α-1-antitrypsin (AAT), a biomarker of epithelial damage in EE *(5)*) (**Fig. 4C**). This suggests that Tuft cells may play a previously unknown role in promoting and responding to inflammation in EE.

To further characterize the EE epithelium, we compared our scRNA-seq dataset to two past studies that used the 10X platform to study intestinal samples. Comparing to a scRNA-seq atlas of the intestine of healthy adults from the U.K., we examined the possibility that small intestinal bacterial overgrowth (SIBO) in EE may lead to a “colonification” of the small intestine due to the increased bacterial load *(57)*. Scoring all epithelial cells in our data for genes differentially upregulated in large vs. small intestine in the healthy human atlas, we did not find evidence for a “colonification” of the small intestine in EE relative to the control cohorts (**Fig. S9A**). In addition, we compared the genes differentially upregulated in EE relative to controls to those found to be upregulated in ulcerative colitis (UC) vs. health in a previous scRNA-seq study *(22)*. This analysis revealed limited overlap between these gene sets, which may highlight key differences in the underlying biology; it may also potentially reflect the difficulty of comparing gene signatures across scRNA-seq technologies (**Fig. S9B**).

### Analysis of the immune compartment in EE reveals increased IFNγ signaling and reduced fractional abundances of T cells expressing markers of tissue resident memory cells

Next, we compared the immune cell compartment between the Zambian cohort with confirmed EE and both the South African and the US cohorts. Cell proportion analysis revealed that, relative to control cohorts, EE samples were enriched for *CD8^hi^* T cells highly expressing the lncRNA *MALAT1* and γδ T cells highly expressing *GZMA* (**Fig. 5A**). This was in agreement with our analyses within the Zambian cohort, where the fractional abundances of these two subsets were significantly associated with disease severity. Consistent with previous findings *(43)*, plasma cells were increased in EE relative to non-EE cohorts despite exhibiting decreased abundances with increased severity within the EE cohort (**Fig. S9C**).

**Figure 5:**
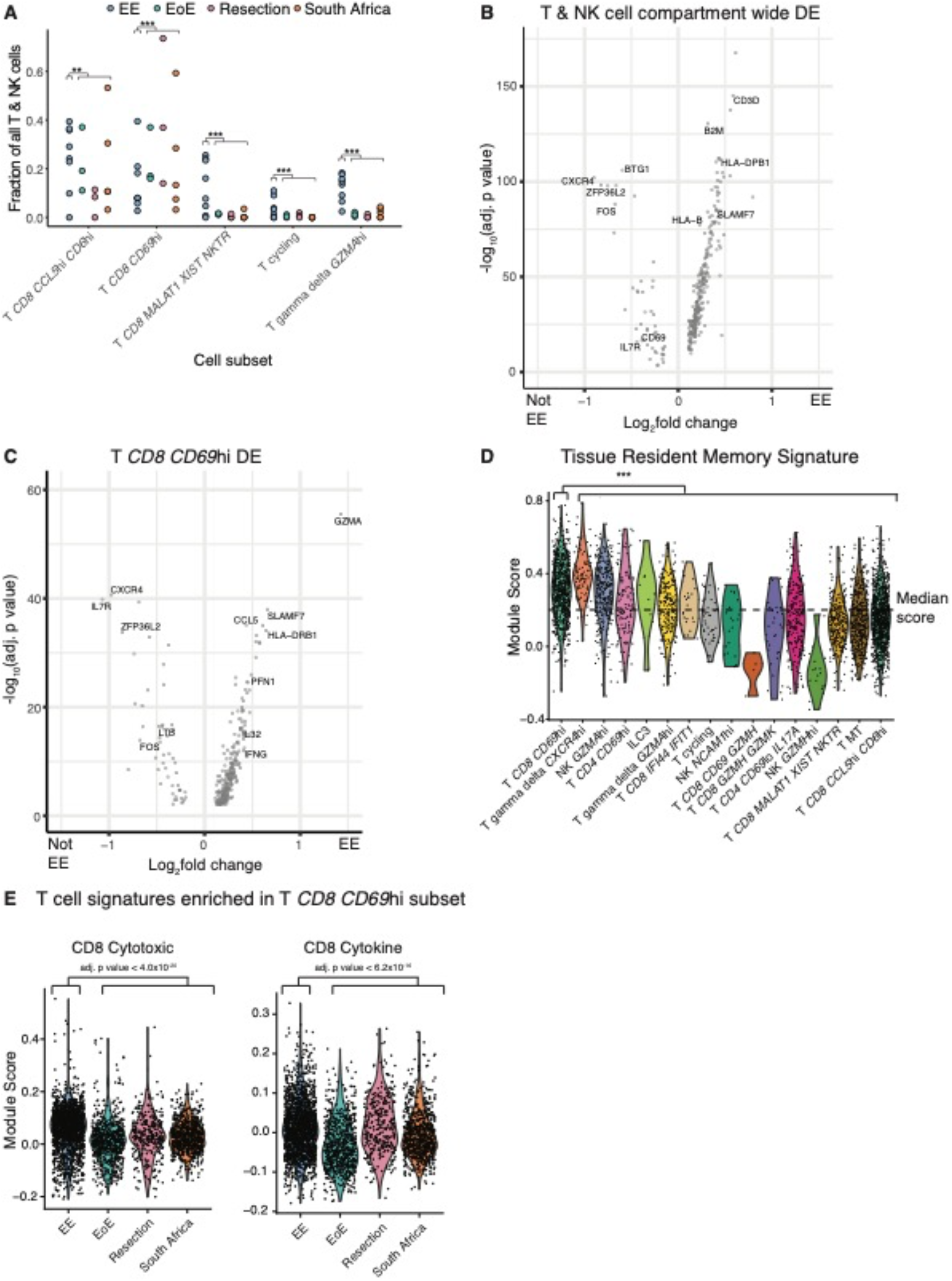
EE is associated with a shift towards activated T cell phenotypes. **A,** Cell subsets with significant shifts in relative abundances between EE and all three control cohorts (*, adj. p< 0.05; **, adj. p < 0.01; ***, adj. p < 0.001; Fischer’s exact test). **B,** Genes differentially expressed in the T & NK cell compartment in EE relative to all three control cohorts. Horizontal and vertical dashed lines respectively refer to an adjusted p value threshold of 0.01 and a log fold change threshold of 0.1. **C,** Genes differentially expressed in in EE relative to all three control cohorts within specific cellular subsets. Horizontal and vertical dashed lines respectively refer to an adjusted p value threshold of 0.01 and a log fold change threshold of 0.1. **D,** Module scores for a tissue resident memory T cell signature in EE relative to all three control cohorts. **E**, T cell activation signatures enriched in T *CD8 CD69*^hi^ cells from EE patients relative to all three control cohorts. Proliferation: adj. p = 3.2*10^−04^, Cohen’s D effect size = 0.32. CD8 Cytotoxic: adj. p = 6.2*10^−25^, Cohen’s D effect size = 0.95. CD8 Cytokine: adj. p = 2.3*10^−23^, Cohen’s D effect size = 0.90

We next identified changes in gene expression in immune cells and found that the majority of gene expression changes in EE relative to the U.S. cohorts occurred within the T cell compartment, rather than the B cell and myeloid compartments (**Fig. 5B, Fig. S10A-C**). In this compartment, genes involved in immune activation *(B2M, HLA-B)* were upregulated whereas downregulated genes included *ZFP36L2* (a negative regulator of T cell proliferation *(58)*), *BTG1* (a factor responsible for T cell quiescence *(59)*), and *IL-7R* (which has been implicated in the formation of memory T cells in response to infection *(60)*) (**Fig. 5B**). Differential expression between EE and other cohorts within each T cell subset revealed that the vast majority of significantly differentially expressed genes within specific cell subsets came from T *CD8 CD69*^hi^ cells, which displayed upregulation of effector-like genes (*IFNG, HLA-DR*, CCL5, IL32, SLAMF7*) in EE and a downregulation of genes (*IL7R* and *CXCR4*) implicated in memory T cell formation in response to infection *(60–62)* (**Fig. 5C, Table S6**). Furthermore, as *CD69* is a potential marker for T cell tissue residency, we sought to ask if the T *CD8 CD69*^hi^ subset represented tissue resident cells. Scoring each T cell subset by a previously published tissue resident memory gene signature, we found that the T *CD8 CD69^hi^* cluster scored highest (adjusted p = 4.98*10^−78^, one sided Wilcoxon test) (**Fig. 5D**) *(63)*. This suggests that this subset consisted of tissue resident cells. To further characterize this subset, we scored all cells from this subset on a set of activation signatures for T cells in scRNA-seq data *(64)*. Relative to the other cohorts, EE T *CD8 CD69*^hi^ cells scored higher for T cells signatures for cytotoxicity and cytokine production (**Fig. 5E**). Together, these results suggested that relative to the other cohorts, the tissue resident T cell niche in EE is more activated and polarized towards production of pro-inflammatory cytokines, which may contribute to the pathogenesis of EE.

To better understand the relationship between our observations in the epithelial and immune compartments in EE, we sought to identify potential ligand-receptor interactions between cellular subsets using NicheNet (**Fig. S11**) *(65)*. In addition to nominating putative drivers of immune epithelial-interactions in EE, this analysis recapitulated our previous observations that relative to the other three cohorts, EE is characterized by increased IFNγ signaling in the epithelium stemming from IFNγ production by T cells, especially the T *CD8 CD69*^hi^ cells. These results further suggest that IFNγ is a critical regulator of EE pathology and may be a valuable avenue for therapeutic approaches to reversing pathogenesis.

### The epithelium from the Zambian EE cohort displays reduced proliferation relative to the U.S. cohorts

Finally, we compared EE samples to those from the U.S. cohorts to identify features of EE that may have been masked in our previous comparisons by any potential features of EE in the South African cohort. Largely, the analyses including and excluding the South African cohort displayed similar results, with the largest difference occurring in the analysis of epithelial cells. Here, differential expression analysis revealed that EE samples displayed compartment-wide downregulation of genes (*KLF4, ATF3, FOS*, and *JUN*) involved in epithelial proliferation *(66)*, IL-22 signaling in the intestine *(67)*, and goblet cell development *(68)* in the confirmed EE samples from Zambia (**Fig. 6A**). Consistently, the confirmed EE samples displayed lower fractional abundances of goblet cells (whose development is induced by IL-22 *(69)*) and ILC3s (a major source of IL-22 in the intestine *(70)*), as well as higher fractional abundances of γδ T cells (which have been shown to reduce IL-22 production by ILC3s in mice fed a low protein diet *(71)*) (**Fig. S8B**). Furthermore, gene set enrichment analysis (GSEA) displayed reduced activity of pro-proliferative MAPK and oxidative phosphorylation signaling in confirmed EE (**Fig. 6B, Table S8**), as well as increased enrichment of the Reactome signature for response of EIF2AAK4 and GCN2 to amino acid deficiency in the epithelial cells of confirmed EE relative to the U.S. cohorts. As amino acid deficiency has been shown to restrict intestinal epithelial proliferation *(72)*, this finding of decreased proliferative signaling and amino acid deficiency response may provide a link connecting low-quality diets in EE endemic areas and the villus regenerative effect of amino acid supplementation in EE populations *(14)*. In addition, goblet cells from patients with confirmed EE upregulated markers for lower crypt goblet cells *(LYZ, REG3A*, and ribosomal genes), suggesting that EE goblet cells have a more immature phenotype and that the lower relative abundances of goblet cells in EE may result from impaired goblet cell maturation *(73)* (**Fig. 6C**). This was consistent with an observed upregulation of *GATA4* in mature enterocytes expressing *APOA* and *ALPI*, as this transcription factor has been shown to bias the intestinal epithelium towards enterocyte differentiation and away from goblet cell differentiation *(74)*. In sum, these findings suggest that EE may be associated with a hypoproliferative state in the epithelium that is accompanied by reduced goblet cell differentiation. This stands in stark contrast to previously characterized inflammation of the small intestine in Crohn’s disease and Celiac disease, where a hyperproliferative epithelium is induced by inflammation *(75, 76)*.

**Figure 6:**
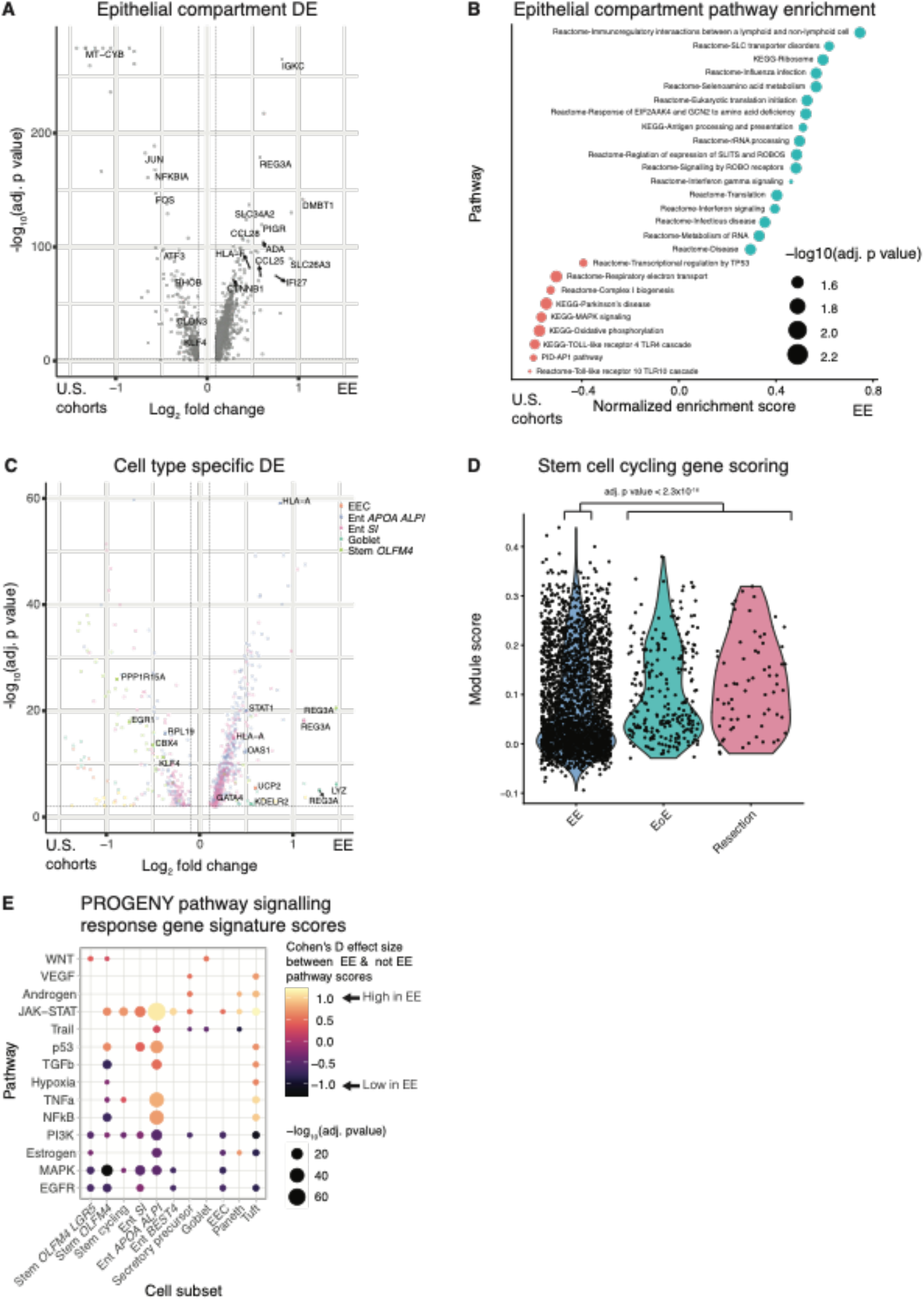
Relative to samples from the U.S. cohorts, the EE epithelium is characterized by decreased proliferative signaling. **A,** Genes differentially expressed in the epithelial compartment in EE relative to the U.S. cohorts. Horizontal and vertical dashed lines respectively refer to an adjusted p value threshold of 0.01 and a log fold change threshold of 0.1. **B,** Gene set enrichment analysis of genes upregulated in epithelial compartment cells in EE relative to the U.S. cohorts. **C,** Genes differentially expressed in EE relative to the U.S. cohorts within specific cellular subsets. Horizontal and vertical dashed lines respectively refer to an adjusted p value threshold of 0.01 and a log fold change threshold of 0.1. **D,** Module scores for cell cycle genes in EE and U.S. cohorts in all cells from stem cell subsets. **E,** PROGENy pathway prediction scores for epithelial cells in EE relative to U.S. cohorts.

To further interrogate potential changes in proliferative capacity, we scored all stem cells in our data on a module of genes annotated as upregulated in cycling human cells *(77)* (which included *MKI67*, cyclins, CDKs, topoisomerases, and polymerases) and found that this module score was significantly reduced in the confirmed EE stem cell compartment compared to the U.S. cohorts (**Fig. 6D**). This further implicated altered proliferation and differentiation in EE relative to the U.S. cohorts. We extended these findings by inferring signaling activity in EE with PROGENy *(53)*, which inferred increased stress responsive pathway activation (JAK/STAT, TNFa, p53) and decreased proliferative pathway activation (EGFR, MAPK, PI3K) in *OLFM4* expressing stem cells (**Fig. 4E, Table S8**). We next inferred upstream transcription factor activity in EE with DoRothEA *(78)*, which revealed reduced activation of ATF2 and ATF4 broadly across the epithelium (**Fig. S12A, Table S9**. This is consistent with reduced IL-22 signaling *(75)* and altered goblet cell development *(79)*). In addition, STAT6, a key regulator of secretory lineage differentiation, was predicted to have lower activity in *OLFM4* expressing stem cells and immature enterocytes *(80)*. This is consistent with predicted upregulation of MYC in secretory precursors and goblet cells, which negatively regulates differentiation towards goblet cells *(81)*. Furthermore, predicted upregulation of GATA4 and GATA6 in *OLFM4* expressing stem cells and mature enterocytes, respectively, suggested that the loss of goblet cell differentiation may occur due to a polarization towards enterocyte differentiation *(74)*.

Altogether these predicted changes indicate altered epithelial differentiation and reduced proliferation of stem cells, the latter of which is surprising as T cell driven enteropathies such as Celiac disease have been shown to induce a hyperproliferative state in the epithelium *(82)*. Thus, we wondered if EE severity within the Zambia cohort would negatively correlate with epithelial proliferation (like in EE relative to the U.S. cohorts) or positive correlate with epithelial proliferation (like in other T cell driven enteropathies). To investigate this question, we correlated PROGENy pathway scores and proliferation gene module scores with histological EE severity scores in the stem cell subsets from HIV-negative patients in the Zambia cohort (**Fig. S1B**). Across all cells from stem cell subsets, the proliferation module weakly positively correlated with EE severity. This suggested that although EE patients as a whole display lower levels of epithelial proliferation relative to U.S. controls, more severe EE leads to relatively higher epithelial proliferation than less severe EE.

## DISCUSSION

Environmental enteropathy is a subclinical condition of the small intestine that contributes to the continuing ill-health and entrenched poverty of millions of people around the world. Here, by utilizing Seq-Well S^3^ – a transformative single-cell transcriptomics platform that can be applied in modestly equipped laboratories – we generated, to our knowledge, the first scRNA-seq dataset profiling EE in the upper small intestine. In the resultant single-cell dataset, we identified a cell subset – surface mucosal cells – which highly expressed *DUOX2* and whose gene expression pattern matched past bulk-transcriptomic signatures of reduced villus height and reduced plasma LPS concentrations in EE. In addition, our dataset revealed regional cell subset fractional abundance differences between the duodenal bulb and distal duodenum, demonstrated that more severe EE is characterized by a shift towards a more intermediate-like epithelial phenotype, provided enhanced resolution of the T cell subsets found in EE, and suggested that HIV enteropathy is associated with distinct cell subsets. In addition, by comparing distal duodenal samples from HIV-negative EE patients to distal duodenal samples from two U.S. cohorts and one South African cohort, we found that EE is characterized by an immature epithelial phenotype and decreased fractional abundance of a T cell subset highly expressing a transcriptional signature of tissue-resident memory cells with increased expression of pro-inflammatory cytokine genes. Furthermore, comparison to only the U.S. cohorts suggested the EE epithelium is characterized by reduced proliferative signaling and fractional abundances of goblet cells. Altogether, our singlecell dataset and analyses re-contextualizes past bulk-transcriptomic studies of EE and identify previously unappreciated correlates of EE pathology.

The presence of surface mucosal cells highly expressing *DUOX2* in EE in our dataset may reflect adaptive remodeling of the epithelium into an intermediate-like wound healing state. One possible means of forming this state may be enterocyte dedifferentiation, which would be consistent with immunohistochemistry staining of EE samples from Bangladeshi children that found DUOX2+ cells at the villus tip *(16)*. While surface mucosal cells have previously been implicated in the mucosal injury response in pyloric metaplasia and in the Ulcer-associated cell lineage (UACL), these conditions are also associated with the presence of mucosal neck cells (high expression of *TFF2* and *MUC6*), whereas mucosal neck cells did not appear to be associated with a wound healing response in our data *(33)*. This may explain why our data suggests surface mucosal cells are a feature of injury response in EE yet past histological analyses of EE did not find any evidence of foveolar metaplasia *(8)*. Furthermore, the presence of this wound healing-like state associated with surface mucosal cells may occur at the expense of absorptive capacity, as absorptive enterocytes are reprogrammed into surface mucosal cells engaged in wound healing. This, in turn, may provide a mechanism that explains why, over time, reduced nutrient absorption in EE is associated with decreased microbial translocation *(83)*. In our study, this phenomenon was clearest in samples from the duodenal bulb from participants with EE, in which we also identified the highest quantity of sequencing reads mapping to the *H. pylori* transcriptome, suggesting that this phenomenon may be due, in part, to extensive *H. pylori* infection. In addition, as *H. pylori* gastritis has been shown often to be associated with duodenal colonization in children *(84)*, *H. pylori* infection may explain why previous studies have found high levels of *DUOX2* transcripts in the distal duodenum of children with EE, whereas our study found *DUOX2* expressing surface mucosal cells in the duodenal bulb and not the distal duodenum of adults with EE. However, this phenomenon may not be unique to EE or to *H. pylori* infection as a previous bulk transcriptomic study has shown a similar tradeoff between *DUOX2* and *APOA4* (a mature enterocyte marker gene) in pediatric Crohn’s disease relative to health *(85)*.

Examination of the other subsets in our dataset provided us with an opportunity to describe variation in EE due to intestinal region, histological severity, and HIV infection. However, this analysis lacked the power to detect significant changes in cell subsets with low fractional abundance and future work is needed to determine if shifts in rarer cell populations are associated with EE severity. In addition, profiling of pediatric patients will be required to understand the generality of these observations and link cellular responses to EE sequalae observed in children. Nonetheless, the data presented here provides a valuable resource for hypothesis generation for future studies of EE.

Comparison of EE to the U.S. and South African cohorts sheds new light on the epithelial and immune cell populations in EE. Increased abundances of immature epithelial cell subsets, increased WNT/ß-catenin signaling, and decreased MAPK signaling suggested that the EE epithelium is biased towards an immature phenotype, which may be a cause or consequence of the villus blunting that is a hallmark of EE pathogenesis. Furthermore, Tuft cells upregulated genes (*ALOX5AP*, *SERPINA1*) involved in promoting and responding to inflammation *(56)*, indicating that this subset may play a previously unappreciated role in inflammation in EE. In addition, we found lower relative abundances of T cells expressing a transcriptional signature of tissue-resident memory T cells and higher relative abundances of plasma cells in EE relative to all other cohorts, suggesting that, similar to recent observations in IBD, EE may be characterized by an imbalance in cell-mediated vs humoral immunity *(86)*. In addition, this tissue-resident memory-like subset highly expressed genes for inflammatory cytokines (including IFNγ), suggesting that although they are present in a lower abundance in EE, these cells may be chronically activated and contribute to pathology. This, in turn, may lead to immune exhaustion and impaired response to new immune stimuli, which may contribute a hindered response to oral vaccines *(87)*. In addition, while IFNγ is often viewed as a pro-inflammatory cytokine in acute inflammation, numerous studies have demonstrated that in chronic inflammation, IFNγ can produce tolerogenic effects *(88)*. While a tolerogenic role for IFNγ would be consistent with reduced oral vaccine efficacy in EE, future work is needed to delineate the specific mechanisms and impacts of IFNγ signaling in the small intestine with EE. Altogether, these identified immune changes provide clues about the mechanisms underlying the lack of oral vaccine efficacy in EE, though future work is needed to corroborate these findings in other cohorts and validate them in animal models.

Our comparison of EE to both the U.S. and South African cohorts may have been confounded by features of moderate EE present in the South African cohort. Thus, we conducted a separate analysis comparing EE to only the U.S. cohorts, and while these results were broadly the same as our previous analysis, the main striking difference was that, relative to the U.S. cohorts, the EE epithelium was characterized by decreased epithelial proliferation, a decreased abundance of ILC3s (a major producer of IL-22 – a positive regulator of intestinal epithelial proliferation *(70)*) and goblet cells (which are induced by IL-22 *(69)*), and increased abundances of γδ T cells (which have been shown to negatively regulate IL-22 production by ILC3s in mice fed a low protein diet *(71)*). Our findings are consistent with past work suggesting decreased proliferation in EE, including a study of fecal transcripts in Malawian children with EE that showed decreased abundances of transcripts in the EGFR and MAPK pathways in the feces of children with EE relative to healthy children *(17)*, as well as mouse studies showing that during *Cryptosporidium* infection protein malnutrition leads to reduced turnover and decreased expression of cell cycle genes in intestinal epithelial cells *(89)*. Consistently, reduced epithelial proliferation in EE relative to the U.S. cohorts was associated with GSEA enrichment of a response to amino acid deficiency. This may explain the reduced relative abundances of goblet cells in EE relative to the U.S. cohorts in our data, as the metabolism of tryptophan – an essential amino acid – has been shown to induce the differentiation of stem cells into goblet cells in mice *(90)*. This is consistent with work showing bulk transcriptomic evidence for decreased tryptophan metabolism in Bangladeshi children with EE *(16)*. Together with recent work showing that amino acid supplementation ameliorates villus blunting in adults with EE *(14)*, our results suggest that amino acid deficiency in the small intestine of adults with EE may lead to hypoproliferative signaling in the epithelium characterized by reduced stem cell proliferation and goblet cell abundance. The fact that these findings are only significant when comparing to the U.S. cohorts but not when including the South African cohorts highlights the need for: 1) follow up mechanistic studies using *in* vitro or animal models to validate these results; and, 2) the difficulty of studying a disease endemic to populations underrepresented in scientific atlases of human health. This calls for the development of diverse single-cell atlases of health that include underrepresented populations to enable better understanding of disease states such as EE.

However, in contrast to the decreased proliferation in EE patients relative to the U.S. cohorts, within HIV-negative Zambian adults with EE, we found that EE histological severity was associated with increased proliferation across all stem cell subsets. This is consistent with past bulk RNA-sequencing of Bangladeshi children with EE, which suggested that more severe EE leads to more proliferation than less severe EE, but lacked the single-cell resolution to demonstrate that this phenomenon was occurring in epithelial stem cells *(16)*. Thus, when comparing EE to U.S. cohorts, EE appears to be negatively associated with stem cell proliferation, whereas when comparing within EE cohorts, more severe EE appears to be positive associated with stem cell proliferation. We believe that this discrepancy is due to the interplay of malnutrition and infection. We propose that the reduced dietary quality (and markedly diminished dietary diversity) in the population of Zambian adults we studied imposes a proliferative constraint on enteropathy. If this is correct, the whole population has reduced stem cell turnover relative to intestinal homeostasis, but some individuals have more proliferation than others in response to infective and inflammatory drivers. However, we acknowledge that dissecting out the different processes induced by reduced dietary quality and pathogen exposure is very difficult, and experimental approaches to this are required. Furthermore, we note that this discrepancy between our within and across country analyses is consistent with past findings in EE research that have found similar differences when performing such comparisons. This includes: 1) a study that found that, relative to a control cohort from the U.K., biopsies from children in the Gambia with EE had higher levels of plasma cells, yet within the Gambian cohort, plasma cell levels decreased with increased malnutrition *(43*); and, 2) work that found that, relative to a control cohort from the UK, Zambian adults have lower levels of expression of the Paneth cell defensin genes *DEFA5* and *DEFA6*, yet within the same cohort of Zambian adults with EE, more severe villus blunting and reduced epithelial surface area correlated with higher levels of defensin gene expression *(12, 91)*. This consistent difference between within and across country analyses of EE illustrates the necessity of comparing to an outgroup to fully understand the pathophysiology of EE.

One intriguing aspect of the observed hypoproliferative state in EE relative to U.S. cohorts is that it stands in stark contrast to the hyperproliferative signaling observed in Crohn’s disease *(75, 92)*. This is of particular interest as limited evidence suggests lower rates of IBD in countries where EE is endemic *(27, 93)*. Thus, we propose that decreased intestinal epithelial proliferation in EE may be protective against inflammation-induced hyperproliferative states in the intestine such as Crohn’s disease. This hypothesis is at an early stage of development and will require validation in other patient cohorts as well as animal or *in vitro* model studies mapping the causal factors driving reduced proliferative signaling. If validated, this result would: 1) nominate genes associated with proliferation in IBD (such as *ATF3*) as potential therapeutic targets for modulating proliferation in EE; and, 2) call for future work investigating whether therapeutics inspired by the causal factors of EE may help to ameliorate IBD.

It is important to recognize that our study has several inherent limitations. We were not able to include a non-EE control group of age-matched adults in Lusaka, Zambia. Thus, we cannot rule out the possibility that the observed differences between EE patients and the U.S. and South African cohorts are due to unobserved variables that differed between these patient populations. In addition, our findings are primarily correlative due to the associative nature of measuring mRNA expression and the difficulties associated with mechanistic follow up validation in humans. This is further limited by the lack of available tissue for histological analysis of samples from the South African cohort. As we saw few stromal cells in our scRNA-seq dataset, our tissue dissociation was likely biased against this subset and future work will be needed to characterize the role of these cells in EE. Future studies of protein deficiency, tryptophan deficiency, and repletion will be necessary for understanding the significance of these results. In addition, the inflamed small intestinal epithelium and lamina propria are highly heterogenous environments containing several rare cell types such as tuft cells, Paneth cells, and a variety of adaptive and innate immune subsets which we did not have sufficient power to analyze in great detail. Furthermore, the HIV status of the participants from the U.S. cohorts was not determined. Additionally, we did not screen EE patients for Celiac disease (which displays a highly similar histological phenotype to EE) and we cannot completely rule out the possibility that some patients may have had Celiac disease, however, we note that in Zambia the staple diet is maize (a gluten free food) and that past studies of EE in Zambian adults have found no evidence of Celiac disease *(94)*. Furthermore, pediatric EE may display different aspects of biology than EE in adults, which calls for future studies in pediatric cohorts. In addition, as EE is an endemic condition in low- and middle-income countries across the globe, it will be necessary to validate that our results hold in cohorts with EE from geographic locations other than Zambia.

Examining our work as a whole, a potential picture of how malnutrition and enteropathogens interact to generate EE pathology emerges. Amino acid deficiency due to a low-quality diet may lead to reduced epithelial proliferation and differentiation towards goblet cells, which in turn would lead to a significant diminution of the host ability to maintain anti-microbial defense of the mucosa and thus may permit increased pathogen-mediated damage of the enterocyte. Increased enteropathogen damage may then lead to epithelial remodeling towards an intermediate wound healing-like phenotype associated with the presence of surface mucosal cells; further work on this intriguing subset is urgently required. In addition, enteropathogen mediated damage would further exaggerate the pathogen-induced IFNγ response and may explain the observed increased IFNγ response in *CD8 CD69^hi^* T cells in our data. Strikingly, metabolites of the essential amino acid tryptophan have been also been shown to induce IL-22 production by ILC3s in the intestine, and thus protein malnutrition may explain the lower relative abundances of ILC3s and the reduced evidence of IL-22 signaling in the epithelium in EE relative to the U.S. cohorts *(95)*. The combined effects of these changes may culminate in reduced IL-22 signaling, thereby promoting a more dormant epithelium, further reducing goblet differentiation, and thereby weakening the host ability to maintain anti-microbial defense. This would represent a vicious cycle of disease that may explain why the effect of combined malnutrition and enteropathogen exposure is far more deleterious than the effect of either factor alone. Together, these findings nominate several therapeutic axes for inducing healthy epithelial proliferation and immune efficacy in patients with EE and potentially other disorders of altered intestinal inflammation and epithelial function like IBD.

## MATERIALS AND METHODS

### Study design

Adult men and women, all volunteers, were recruited from a disadvantaged community in Lusaka, Zambia, in which we have carried out previous studies of environmental enteropathy *(11, 15)*. All volunteers gave written, fully informed consent. The study was approved by the University of Zambia Biomedical Research Ethics Committee (reference 006-11-15, dated 12^th^ January 2016). From July 2018 to August 2018, volunteers underwent endoscopy with a Pentax 2990i gastroscope or a Pentax VSB2990i enteroscope, in the endoscopy unit of the University Teaching Hospital, Lusaka, under sedation with diazepam and pethidine. Duodenal tissue was collected from eosinophilic esophagitis (EoE) patients undergoing surveillance gastroscopy at Massachusetts General Hospital (MGH), Boston. Informed consent was obtained from EoE patients under a protocol approved by MGH. Resection samples were obtained from patients undergoing duodenal resection for pancreatic cancer (but in whom no local spread was apparent) in accordance with MGH IRB guidance under Mass General Brigham Protocol 2010P000632. Informed consent was obtained from participants recruited into this study at the Inkosi Albert Luthuli Central Hospital in Durban, South African.

### Biopsy handling and tissue digestion

Two biopsies from the patients with environmental enteropathy were collected into normal saline, then orientated under a dissecting microscope, fixed in buffered formal saline, and processed to 3μm sections for haematoxylin/eosin staining. These sections were scanned using an Olympus VS120 scanning microscope, measured for average villus height (VH) and crypt depth (CD), and scored using a new scoring system for environmental enteropathy, which captures information about villus and crypt architecture, lamina propria and epithelial inflammation, epithelial injury and detachment, and abnormalities of Paneth and goblet cells *(8)*. Biopsies from EE, EoE, resection, and South African patients were dissociated into single-cell suspensions using a modified version of a previously published protocol *(22)*. Further detail for tissue digestion is provided in the supplementary methods.

### Single-cell RNA-sequencing with Seq-Well S^3^

Please refer to the supplementary methods for further detail. Briefly, the epithelial and lamina propria layers of the biopsies were dissociated into single-cell suspensions. Then, ~16,000 singlecells (~8,000 epithelial and ~8,000 lamina propria) were loaded onto a functionalized polydimethylsiloxane (PDMS) array preloaded with ~80,000 uniquely-barcoded mRNA capture beads (Chemgenes; MACOSKO-2011-10), and sequencing libraries were obtained following the protocol described in Hughes et al. *(20)* Arrays were sequenced on an Illumina Next-Seq. Sequencing read alignment and demultiplexing was performed on the cumulus platform *(96)* using snapshot 6 of the Drop-seq pipeline previously described in *(77)*, resulting in a cell barcode by UMI digital gene expression (DGE) matrix. QC was performed to remove low quality cell barcodes and doublet cells. To identify cell subsets, we adopted an existing pipeline for automated iterative clustering of single-cell data (pipeline described in detail in supplementary methods) that has been shown to identify batch effects without collapsing distinct rare cell types *(28, 29, 97)*. We applied this pipeline to the data from the EE and U.S. cohorts. To facilitate visualization and presentation of the data, final cell subsets obtained from automated iterative clustering and manual annotation with lineage markers were grouped into 1) epithelial 2) T & NK cells and 3) B cells, myeloid cells, and stromal cells. To correct for possible batch effects between data collected in different laboratories while allowing for the possibility of new cell type discovery, we integrated our data with the South African dataset with the Seurat V3 method *(98)*. Cells from each group were then normalized with SCTransform and PCA and UMAP embeddings were calculated for visualization. Furthermore, final cell types were input to the FindAllMarkers function in Seurat with default parameters and a Wilcoxon test, producing marker genes for each final cell type.

All subsets were scored on gene signatures of reduced villus height and decreased and plasma LPS signatures from Chama et al. *(7)* using the AddModuleScore function in Seurat and a Wilcoxon test was used to assess the significance of the difference of the mean module scores in surface mucosal cells relative to all other subsets. Epithelial trajectories were inferred with PAGA *(35)* in Scanpy *(99)* run with default parameters. Prior to analysis in Scanpy, data was normalized in Seurat with SCTransform. Prior to running PAGA, the nearest neighbor graph was calculated in Scanpy with neighbors = ceiling(0.5(number of epithelial cells)^0.5)). RNA velocity trajectories were inferred with velocyto using default parameters *(36)*.

### Analyses examining variation within samples from the EE cohort

To assess the epithelial subsets associated with surface mucosal cells in duodenal bulb samples from EE patients, we calculated the Pearson correlation between the fractional abundances of all epithelial subsets within these subsets. We then hierarchically clustered the resulting correlations using the k-means approach implemented in the ComplexHeatMap package in R.

Changes in the relative abundances of cell subsets by differing HIV infection status and small intestinal region were detected by a leave-one-out approach in order to avoid identifying patient specific effects. For each comparison (HIV positive vs negative, intestinal region A vs other intestinal regions, etc.), across all patients, a Fisher’s exact test was used to identify subsets with significant changes in relative abundance (adjusted p value < 0.05). Then this analysis was repeated with each patient left out and only cell subsets that were significant in all analyses with patients left out were included in the final results.

Cell subsets significantly associated with histological severity from the Zambian cohort were identified by running Dirichlet Regression, which allows for testing for differences in cell subsets along a continuous dependent variable, with the R package DirichletReg. Further details are provided in the supplementary methods.

### Comparison of distal duodenal samples from HIV-negative EE and U.S. control cohorts

Distal duodenal samples from patients with HIV-negative EE were compared to distal duodenal samples from HIV-negative EE patients to distal duodenal samples from patients with EoE and from tissue resections from patients with pancreatic cancer. In all analyses for this comparison, we sought to identify biological features driven by variation in EE biology relative to the two control cohorts (as opposed to identifying biology that distinguished only one cohort from EE). Thus, in all subsequent analyses for this comparison we required that results pass the following two criteria: 1) Result significant when comparing EE vs both control cohorts 2) Result displayed the same direction of change between EE and each control cohort. Following this approach, we conducted compositional testing, differential expression, PROGENy, DoRothEA, GSEA, NicheNet, and comparison with gene signatures from existing scRNA-seq dataset analyses. Further details for each of these analyses in provided in the supplementary methods.

### Metagenomic mapping of *H. pylori* reads with Kraken2

Bam files from sequencing were classified with Kraken2 with the following pipeline from the Broad institute: https://dockstore.org/workflows/github.com/broadinstitute/viral-pipelines/classify_kraken2:master?tab=files. For more detail, see the supplementary methods. *(100)*

### Immunohistochemical staining

Formalin fixed paraffin embedded slides were antigen retrieved using heat induced epitope retrieval at 97 degrees C for 20 minues with citrate buffer pH 6. Staining was carred out on a ThermoScientific ICH Autostainer 360 with a 10 minute endogenous peroxidase blocking step, a 30 minute protein blocking step, a 60 minute primary antibody step (DUOX2 Millipore Sigma MABN787 at a 1:200 dilution), a 15 minute labeled polymer step, and a 5 minute DAB step.

## Supplementary Materials

Materials and Methods

Fig. S1: Characterization of epithelial cell subsets revealed by single-cell RNA-sequencing.

Fig. S2: Characterization of T and NK cell subsets revealed by single-cell RNA-sequencing.

Fig. S3: Characterization of B cell, myeloid, and stromal subsets revealed by single-cell RNA-sequencing.

Fig. S4: Number of genes and UMIs per cell across samples

Fig. S5: Characterization of surface mucosal and dedifferentiation-like subsets

Fig. S6: Images of immunohistochemical staining of DUOX2 protein

Fig. S7: Variation in EE biology associated with HIV infection

Fig. S8: Samples from South African participants display features of EE

Fig. S9: Comparing EE epithelial cells to signatures from existing scRNA-seq datasets of the human gastrointestinal tract

Fig. S10: Cell subset relative abundance and differential expression changes in B cells and myeloid cells between EE and controls

Fig. S11: NicheNet analysis of cell-cell signaling in EE relative to controls

Fig. S12: Additional characterization of epithelial signaling changes in the EE cohort relative to the U.S. cohorts

Table S1: Clinical characteristics of donors of intestinal biopsy samples

Table S2: Patient cohort, intestinal region, and HIV infection status of participants studied

Table S3: Villus morphometry for EE distal duodenal samples

Table S4: Marker genes for all subsets

Table S5: Histological severity scores for EE biopsies with matched histology

Table S6: Genes differentially expressed in 1) HIV-EE distal duodenal samples relative to samples from U.S & South African cohorts 2) HIV-EE distal duodenal samples relative to samples from U.S cohorts

Table S7: KEGG, REACTOME, and PID pathways differentially enriched in 1) HIV-EE distal duodenal samples relative to samples from U.S & South African cohorts 2) HIV-EE distal duodenal samples relative to samples from U.S cohorts

Table S8: PROGENY pathway activation scores differentially regulated in epithelial cells in 1) HIV-EE distal duodenal samples relative to samples from U.S & South African cohorts 2) HIV-EE distal duodenal samples relative to samples from U.S cohorts

Table S9: Transcription factors with upstream activity predictions from DoRothEA differentially scored in epithelial cells in 1) HIV-EE distal duodenal samples relative to samples from U.S & South African cohorts 2) HIV-EE distal duodenal samples relative to samples from U.S cohorts

## Acknowledgements

We thank the participants who participated in this study; B. Mead, S. Nguyen, A. Rubin, and R. Xavier for insightful feedback and copy editing; Chola Mulenga for histological processing and morphometry; Rose Banda for recruitment of the Zambian volunteers and sample collection; and Joyce Sibwani and Rose Soko for expert endoscopy nursing. Figure 1b and the graphical abstract were generated with Biorender.com.

## Funding

This work was supported, in part, by grants to PK from Barts & The London Charity (G-000907) and from the Bill & Melinda Gates Foundation (OPP1066118). A.K.S. was supported was supported, in part, by the Searle Scholars Program, the Beckman Young Investigator Program, Sloan Fellowship in Chemistry, the NIH (5U24AI118672), the Bill and Melinda Gates Foundation, and the Ragon Institute. T.W. was supported by the NIH (DK007762). J.O.M is a New York Stem Cell Foundation – Robertson Investigator. J.O.M is a New York Stem Cell Foundation – Robertson Investigator. J.O.M was supported by the Richard and Susan Smith Family Foundation, the HHMI Damon Runyon Cancer Research Foundation Fellowship (DRG-2274-16), the AGA Research Foundation’s AGA-Takeda Pharmaceuticals Research Scholar Award in IBD – AGA2020-13-01, the HDDC Pilot and Feasibility P30 DK034854, the Food Allergy Science Initiative, the Leona M. and Harry B. Helmsley Charitable Trust, The Pew Charitable Trusts Biomedical Scholars and The New York Stem Cell Foundation. T.K.H. is supported by the NIH (F30-AI143160-01A1).

## Author Contributions

C.K., T.W. T.K.H., Z.G., A.K.S., and P.K. conceived and designed experiments; C.K. performed tissue dissociation with assistance from T.K.H. J.O.M., and M.V.; C.K., T.K.H., T.W., S.M., I.F., and M.V. performed scRNA-seq; C.K. and N.M. analyzed the single-cell data; B.D. and J.O.M. developed code for iterative tiered clustering; V.M., E.B., and K.Z. performed histological analyses; J.T.B. coordinated duodenal resection samples; A.M.U. and J.J.G. coordinated distal duodenal samples from patients with EoE; C.K., T.W., T.K.H., J.O.M., Z.G., A.K.S., and P.K. interpreted the data; C.K., T.W., A.K.S., and P.K. wrote the manuscript with input from all authors.

## Competing interests

J.O.M. reports compensation for consulting services with Cellarity and Hovione. A.K.S. reports compensation for consulting and/or SAB membership from Merck, Honeycomb Biotechnologies, Cellarity, Repertoire Immune Medicines, Hovione, Third Rock Ventures, Ochre Bio, FL82, Relation Therapeutics, and Dahlia Biosciences unrelated to this work. A.K.S. has received research support from Merck, Novartis, Leo Pharma, Janssen, the Bill and Melinda Gates Foundation, the Moore Foundation, the Pew-Stewart Trust, Foundation MIT, the Chan Zuckerberg Initiative, Novo Nordisk and the FDA unrelated to this work. T.K.H. has served as a consultant and holds equity in nference, inc. The remaining authors disclose no conflicts.

## Data availability

For all samples, the expression matrix and associated metadata can be visualized and downloaded from the Alexandria Project, a Bill and Melinda Gates Foundation-funded portal (part of the Single-Cell Portal hosted by The Broad Institute of MIT and Harvard): https://singlecell.broadinstitute.org/single_cell/study/SCP1307.

## Abbreviations

(CD): Crypt depth
(EE): Environmental enteropathy
(EoE): Eosinophilic esophagitis
(SAM): Severe acute malnutrition
(scRNA-seq): single-cell RNA-sequencing
(VH): Villus height

## Notes

https://singlecell.broadinstitute.org/single_cell/study/SCP1307

