## Supplemental Figures and Methods for "Single-cell profiling of environmental enteropathy reveals signatures of epithelial remodeling and immune activation in severe disease"

#### SUPPLEMENTARY MATERIALS

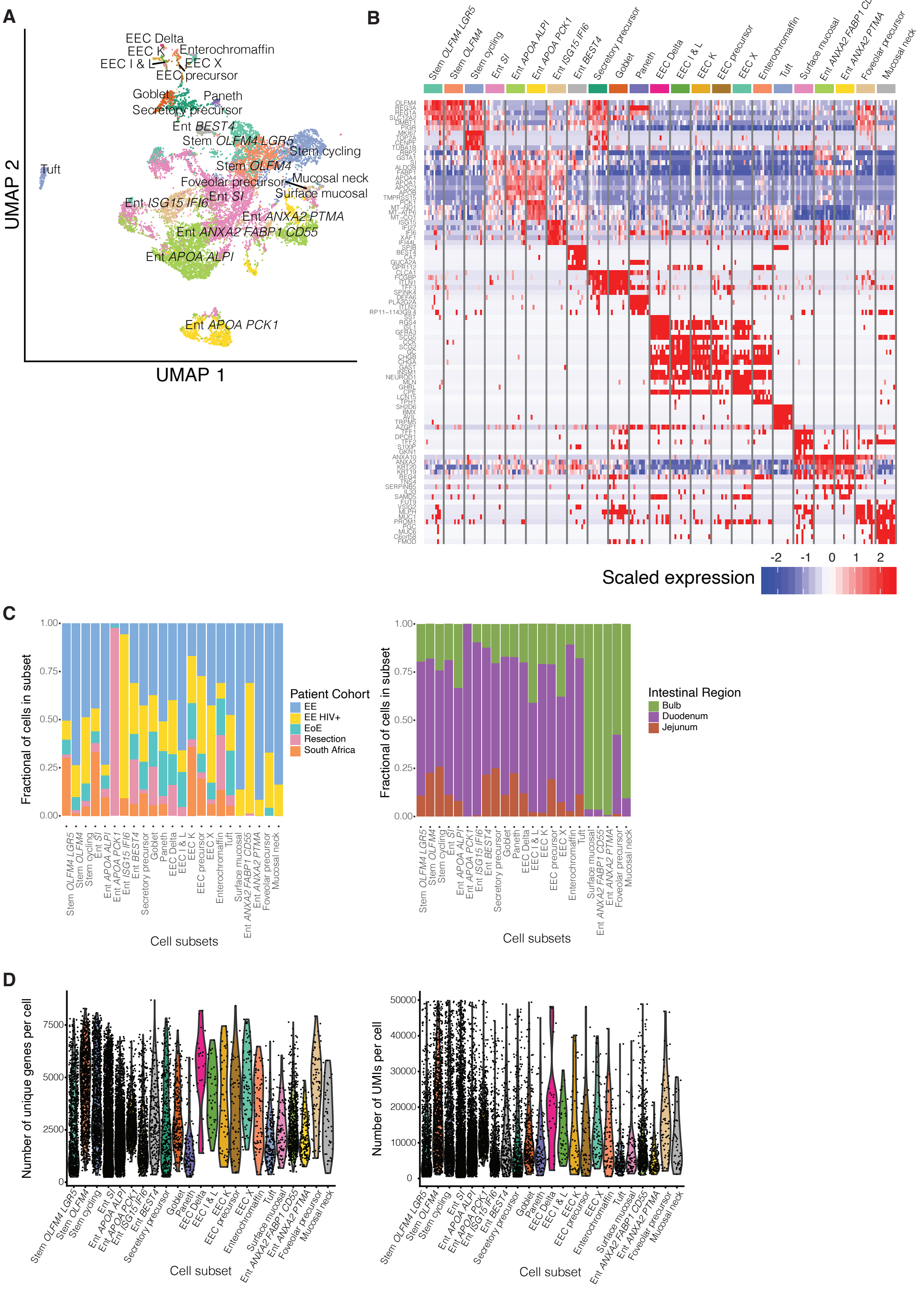

A
EEC Delta
EEC K
Enterochromaffin
EEC I & L
EEC X
EEC precursor
Goblet
Paneth
Secretory precursor
Ent BEST4
Stem OLFM4 LGR5
Stem cycling
Tuft
Stem OLFM4
Foveolar precursor
Mucosal neck
Ent SI
Surface mucosal
Ent ISG15 IFI6
Ent ANXA2 PTMA
Ent ANXA2 FABP1 CD55
Ent APOA ALPI
Ent APOA PCK1
UMAP 2
UMAP 1
B
Stem OLFM4 LGR5
Stem OLFM4
Stem cycling
Ent SI
Ent APOA ALPI
Ent APOA PCK1
Ent ISG15 IFI6
Ent BEST4
Secretory precursor
Goblet
Paneth
EEC Delta
EEC I & L
EEC K
EEC precursor
EEC X
Enterochromaffin
Tuft
Surface mucosal
Ent ANXA2 FABP1 CD55
Ent ANXA2 PTMA
Foveolar precursor
Mucosal neck
Scaled expression
-2 -1 0 1 2
C
Fractional of cells in subset
1.00
0.75
0.50
0.25
0.00
Patient Cohort
EE
EE HIV+
EoE
Resection
South Africa
Cell subsets
Intestinal Region
Bulb
Duodenum
Jejunum
D
Number of unique genes per cell
7500
5000
2500
0
Cell subset
Number of UMIs per cell
50000
40000
30000
20000
10000
0
Cell subset

**Figure S1: Characterization of epithelial cell subsets revealed by scRNA-seq analysis.**

**A,** UMAP visualization of epithelial subsets

**B,** Heatmap of the top five marker genes (Wilcoxon test) of each epithelial subset

**C,** Fraction of cells in each subset from each patient cohort; from HIV-positive vs. HIV-negative patients; and from each intestinal region.

**D**, Violin plots by cell subset of the number of unique molecular identifiers (UMIs) per cell and the number of unique genes detected per cell.

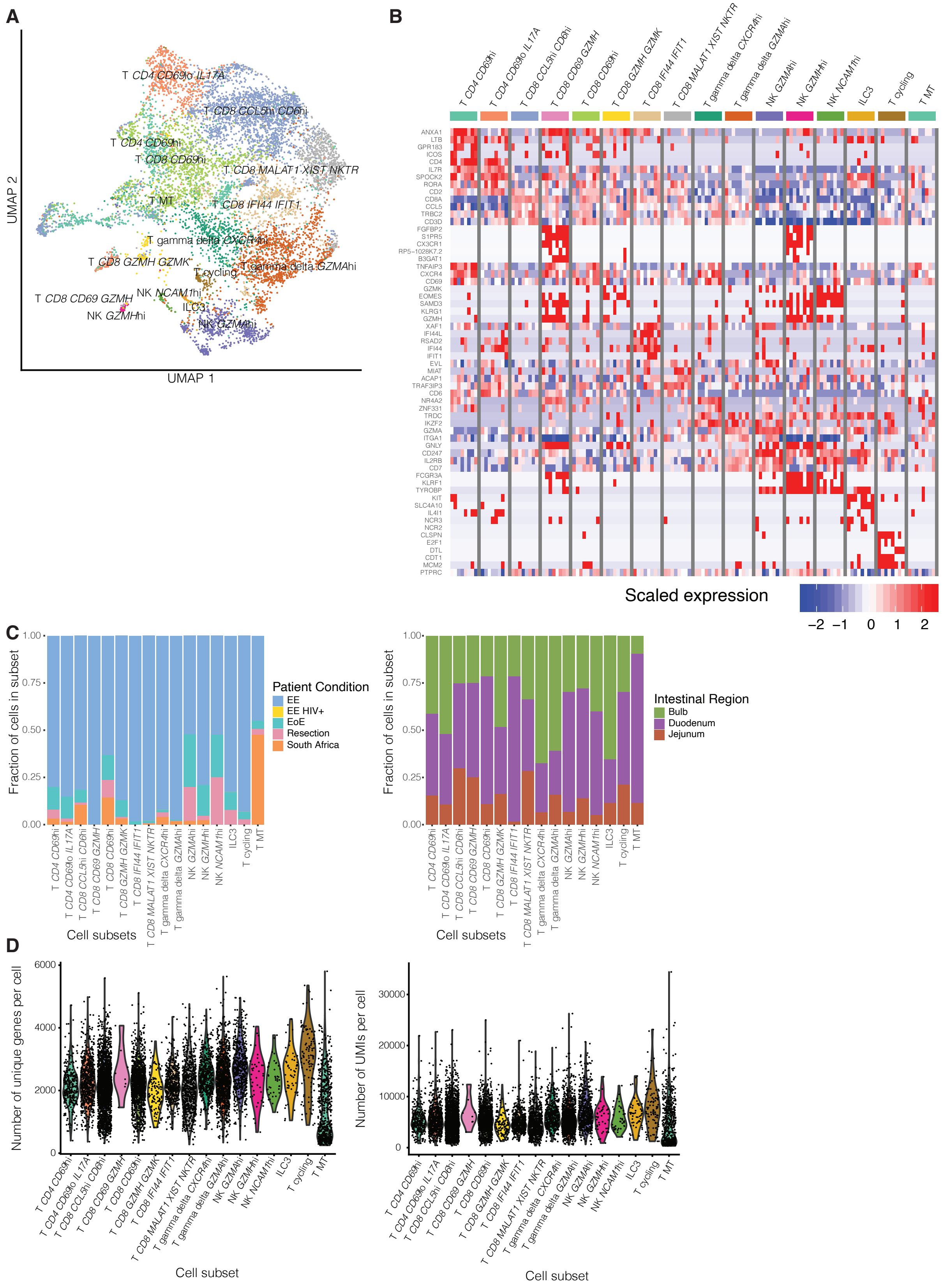

A
UMAP 2
UMAP 1
T CD4 CD69lo IL17A
T CD8 CCL5hi CD6hi
T CD4 CD69hi
T CD8 CD69hi
T CD8 MALAT1 XIST NKTR
T MT
T CD8 IFI44 IFIT1
T gamma delta CXCR4hi
T CD8 GZMH GZMK
T cycling
T gamma delta GZMAhi
T CD8 CD69 GZMH
NK NCAM1hi
NK GZMHhi
ILC3
NK GZMAhi
B
T CD4 CD69hi
T CD4 CD69lo IL17A
T CD8 CCL5hi CD6hi
T CD8 CD69 GZMH
T CD8 CD69hi
T CD8 GZMH GZMK
T CD8 IFI44 IFIT1
T CD8 MALAT1 XIST NKTR
T gamma delta CXCR4hi
T gamma delta GZMAhi
NK GZMAhi
NK GZMHhi
NK NCAM1hi
ILC3
T cycling
T MT
ANXA1
LTB
GPR183
ICOS
CD4
IL7R
SPOCK2
RORA
CD2
CD8A
CCL5
TRBC2
CD3D
FGFBP2
S1PR5
CX3CR1
RP5−1028K7.2
B3GAT1
TNFAIP3
CXCR4
CD69
GZMK
EOMES
SAMD3
KLRG1
GZMH
XAF1
IFI44L
RSAD2
IFI44
IFIT1
EVL
MIAT
ACAP1
TRAF3IP3
CD6
NR4A2
ZNF331
TRDC
IKZF2
GZMA
ITGA1
GNLY
CD247
IL2RB
CD7
FCGR3A
KLRF1
TYROBP
KIT
SLC4A10
IL4I1
NCR3
NCR2
CLSPN
E2F1
DTL
CDT1
MCM2
PTPRC
Scaled expression
−2 −1 0 1 2
C
Fraction of cells in subset
1.00 0.75 0.50 0.25 0.00
Patient Condition
EE
EE HIV+
EoE
Resection
South Africa
Cell subsets
Intestinal Region
Bulb
Duodenum
Jejunum
Cell subsets
D
Number of unique genes per cell
6000 4000 2000 0
Cell subset
Number of UMIs per cell
30000 20000 10000 0
Cell subset

**Figure S2: Characterization of T and NK cell subsets revealed by scRNA-seq analysis.**

**A,** UMAP visualization of epithelial subsets

**B,** Heatmap of the top five marker genes (Wilcoxon test) of each T and NK cell subset

**C,** Fraction of cells in each subset from each patient cohort; from HIV-positive vs. HIV-negative patients; and from each intestinal region.

**D**, Violin plots by cell subset of the number of unique molecular identifiers (UMIs) per cell and the number of unique genes detected per cell.

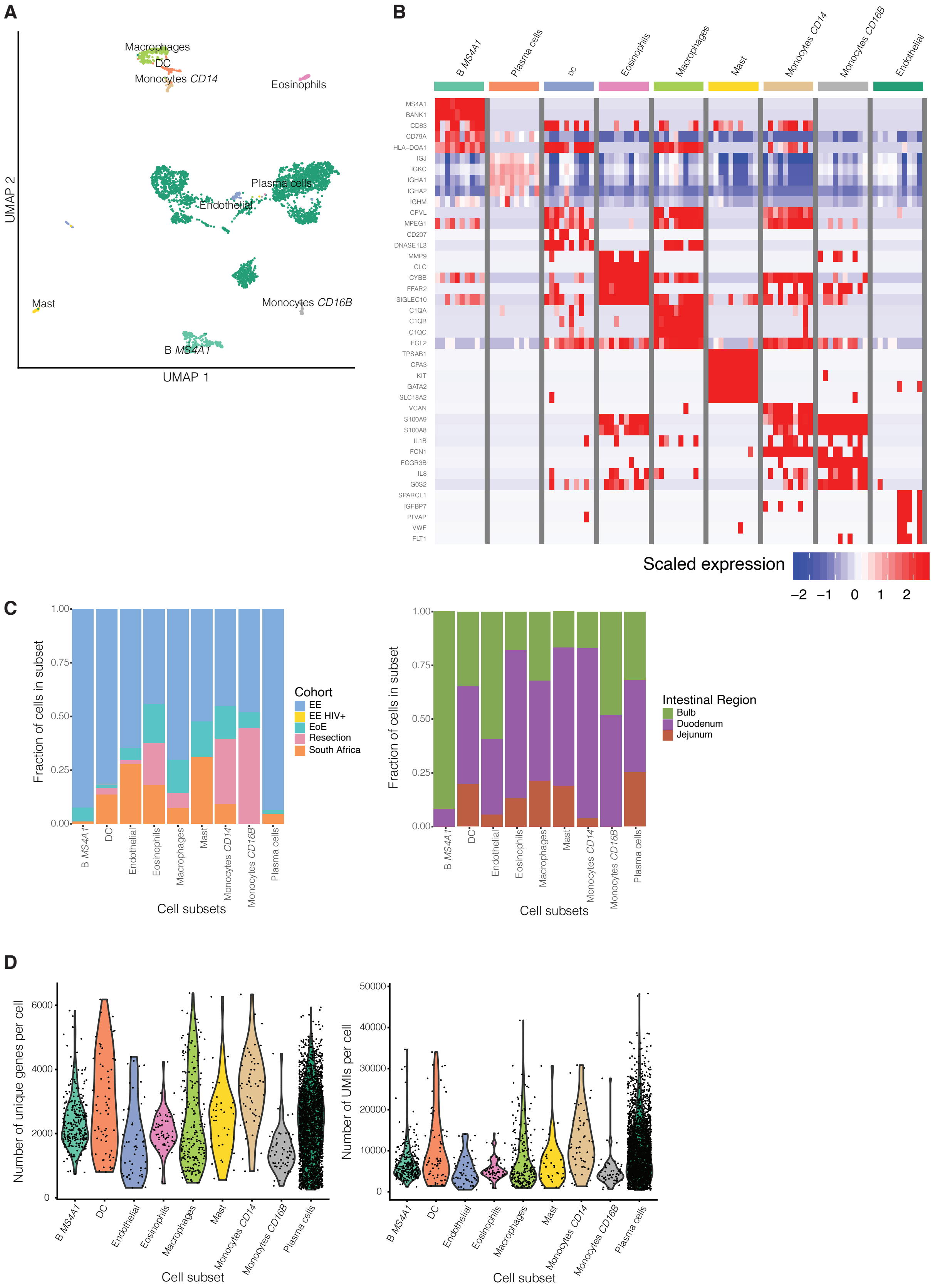
A
Macrophages
DC
Monocytes CD14
Eosinophils
Plasma cells
Endothelial
Mast
Monocytes CD16B
B MS4A1
UMAP 2
UMAP 1
B
B MS4A1
Plasma cells
DC
Eosinophils
Macrophages
Mast
Monocytes CD14
Monocytes CD16B
Endothelial
MS4A1
BANK1
CD83
CD79A
HLA-DQA1
IGJ
IGKC
IGHA1
IGHA2
IGHM
CPVL
MPEG1
CD207
DNASE1L3
MMP9
CLC
CYBB
FFAR2
SIGLEC10
C1QA
C1QB
C1QC
FGL2
TPSAB1
CPA3
KIT
GATA2
SLC18A2
VCAN
S100A9
S100A8
IL1B
FCN1
FCGR3B
IL8
G0S2
SPARCL1
IGFBP7
PLVAP
VWF
FLT1
Scaled expression
−2 −1 0 1 2
C
Fraction of cells in subset
0.00
0.25
0.50
0.75
1.00
Cohort
EE
EE HIV+
EoE
Resection
South Africa
Intestinal Region
Bulb
Duodenum
Jejunum
B MS4A1
DC
Endothelial
Eosinophils
Macrophages
Mast
Monocytes CD14
Monocytes CD16B
Plasma cells
Cell subsets
D
Number of unique genes per cell
0
2000
4000
6000
Number of UMIs per cell
0
10000
20000
30000
40000
50000
Cell subset

**Figure S3: Characterization of B cell, myeloid, and stromal subsets revealed by scRNA-seq analysis.**

**A,** UMAP visualization of epithelial subsets

**B,** Heatmap of the top five marker genes (Wilcoxon test) of each B cell, myeloid, and stromal cell subset

**C,** Fraction of cells in each subset from each patient cohort; from HIV-positive vs. HIV-negative patients; and from each intestinal region.

**D**, Violin plots by cell subset of the number of unique molecular identifiers (UMIs) per cell and the number of unique genes detected per cell.

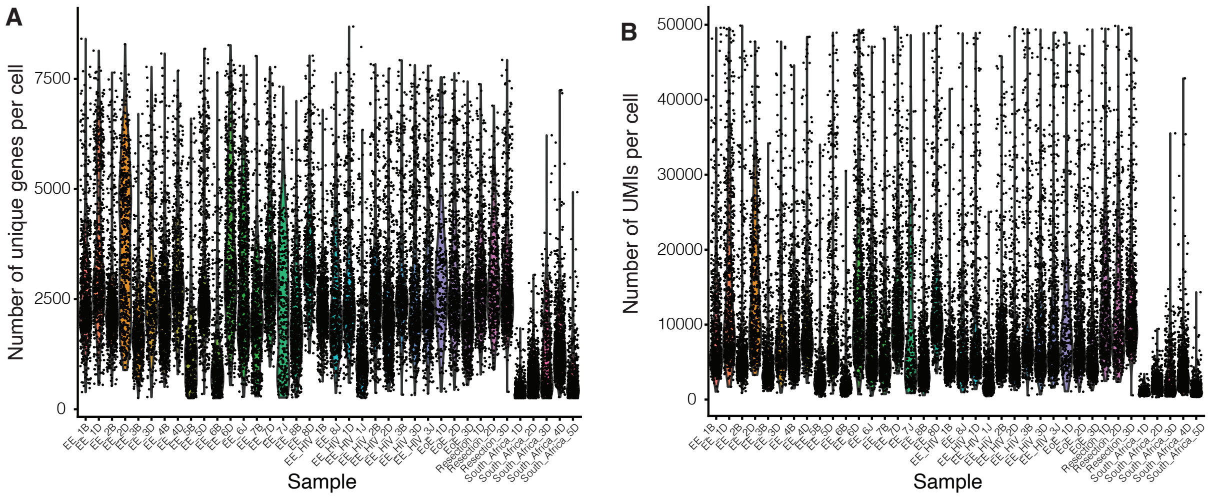
A
Number of unique genes per cell
0
2500
5000
7500
Sample
B
Number of UMIs per cell
0
10000
20000
30000
40000
50000
Sample

**Figure S4: Number of genes and UMIs per cell across samples**

**A,** Violin plot of number of genes per cell across samples

**B,** Violin plot of number of UMIs per cell across samples

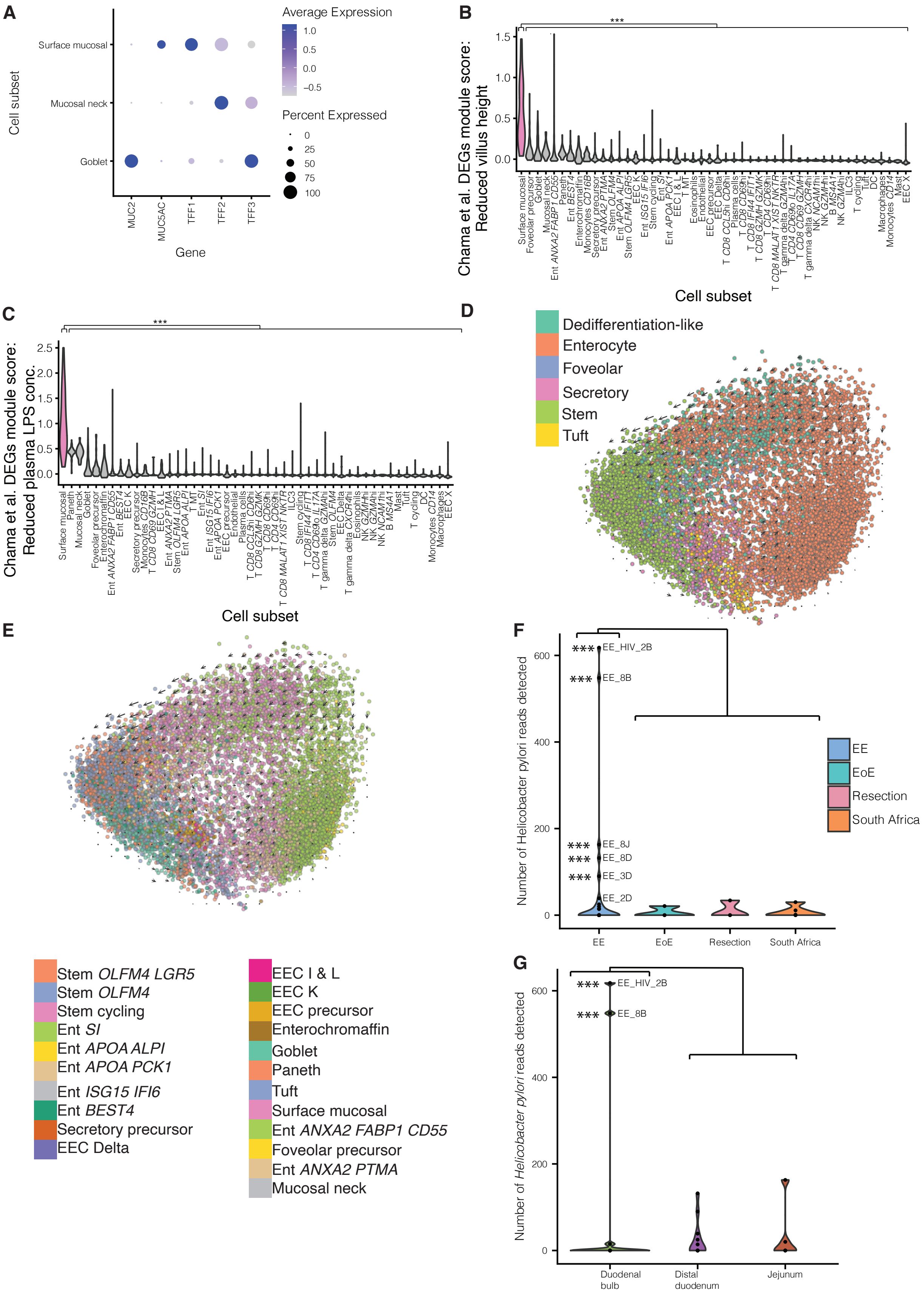

A
Average Expression
1.0
0.5
0.0
−0.5
Percent Expressed
0
25
50
75
100
Cell subset
Surface mucosal
Mucosal neck
Goblet
MUC2
MUC5AC
TFF1
TFF2
TFF3
Gene
B
***
Chama et al. DEGs module score:
Reduced villus height
Cell subset
C
***
Chama et al. DEGs module score:
Reduced plasma LPS conc.
Cell subset
D
Dedifferentiation-like
Enterocyte
Foveolar
Secretory
Stem
Tuft
E
Stem OLFM4 LGR5
Stem OLFM4
Stem cycling
Ent SI
Ent APOA ALPI
Ent APOA PCK1
Ent ISG15 IFI6
Ent BEST4
Secretory precursor
EEC Delta
EEC I & L
EEC K
EEC precursor
Enterochromaffin
Goblet
Paneth
Tuft
Surface mucosal
Ent ANXA2 FABP1 CD55
Foveolar precursor
Ent ANXA2 PTMA
Mucosal neck
F
Number of Helicobacter pylori reads detected
*** EE_HIV_2B
*** EE_8B
*** EE_8J
*** EE_8D
*** EE_3D
EE_2D
EE
EoE
Resection
South Africa
G
Number of Helicobacter pylori reads detected
*** EE_HIV_2B
*** EE_8B
Duodenal bulb
Distal duodenum
Jejunum

**Figure S5: Characterization of surface mucosal and dedifferentiation-like subsets**

**A,** Expression of mucin and trefoil factor genes distinguishing Surface mucosal, mucosal neck, and goblet cell subsets. Dot size represents the fraction of a cell subset (rows) expressing a given gene (columns). Dot hue represents the scaled average expression by gene column.

**B,** Violin plot scoring all subsets on a module score generated from genes differentially upregulated in bulk RNA-sequencing of samples with EE and reduced villus height (VH) in Chama et al. Surface mucosal cells were enriched for this signature relative to all other subsets (***, $p < 0.001$; Wilcoxon test)

**C,** Violin plot scoring all subsets on a module score generated from genes differentially upregulated in bulk RNA-sequencing of samples with EE and decreased plasma LPS concentrations in Chama et al. Surface mucosal cells were enriched for this signature relative to all other subsets (***, $p < 0.001$, Wilcoxon test)

**D**, Velocyto results grouped by epithelial differentiation trajectory identified in Figure 2e

**E,** Velocyto results grouped by epithelial subset

**F,** Violin plots by participant cohort of the number of *H. pylori* reads detected by metagenomic alignment with Kraken2 in the sequencing reads from each sample (T-test: *,adj. $p < 0.05$; **, adj. $p < 0.01$; ***, adj. $p < 0.001$)..

**G,** Violin plots by intestinal region of the number of *H. pylori* reads detected by metagenomic alignment with Kraken2 in the sequencing reads from each sample from participants in the Zambian EE cohort (T-test: *,adj. $p < 0.05$; **, adj. $p < 0.01$; ***, adj. $p < 0.001$).

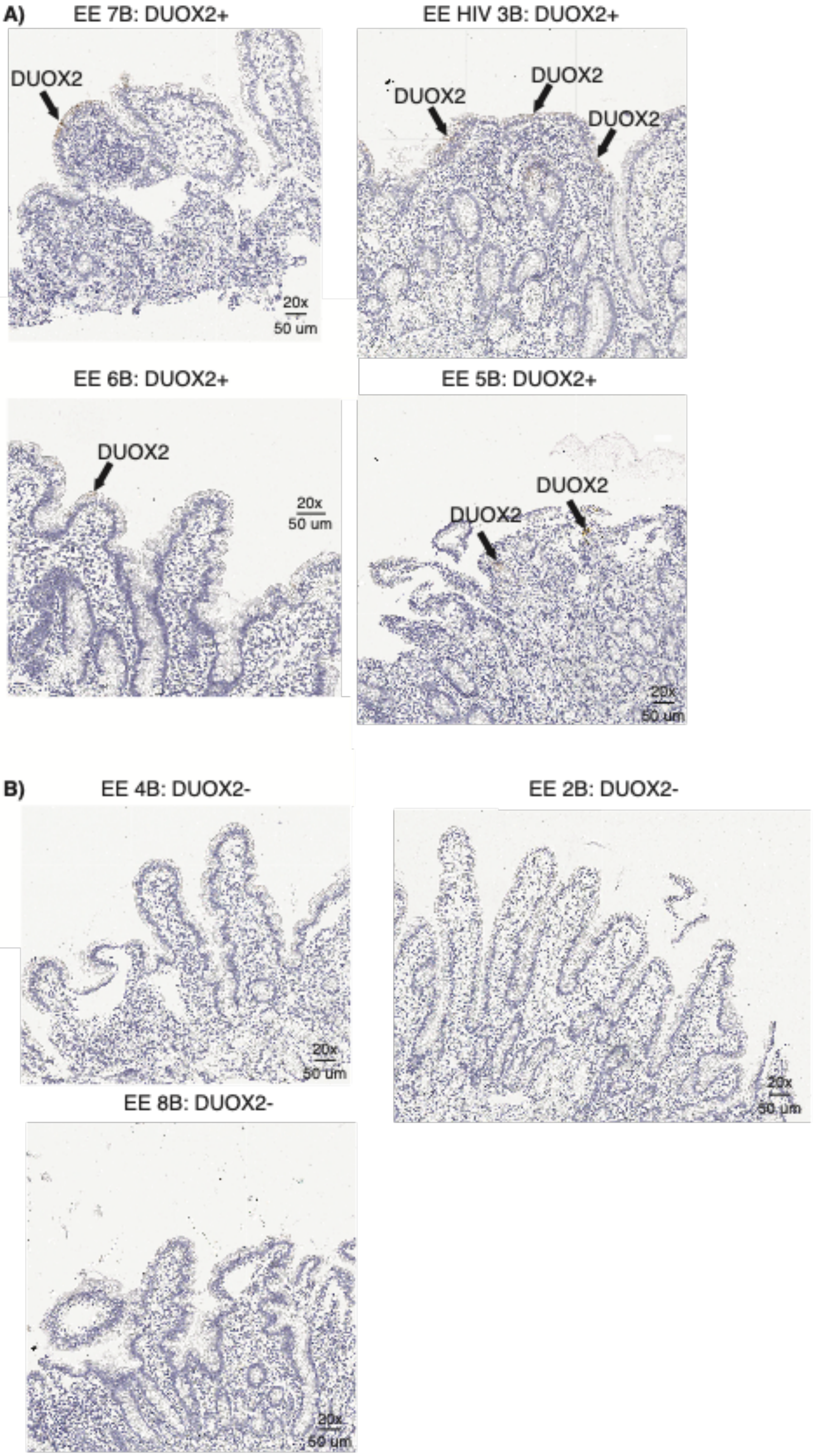

A)
EE 7B: DUOX2+
DUOX2
20x
50 um
EE HIV 3B: DUOX2+
DUOX2
DUOX2
DUOX2
EE 6B: DUOX2+
DUOX2
20x
50 um
EE 5B: DUOX2+
DUOX2
DUOX2
20x
50 um
B)
EE 4B: DUOX2-
20x
50 um
EE 2B: DUOX2-
20x
50 um
EE 8B: DUOX2-
20x
50 um

**Figure S6: Images of immunohistochemical staining of DUOX2 protein**

**A,** H&E images (purple, H&E) that stained positive for DUOX2 (brown, DUOX2) from duodenal bulb samples from participants in the Zambian EE cohort

**B,** H&E images (purple, H&E) that did not stain positive for DUOX2 (brown, DUOX2) from duodenal bulb samples from participants in the Zambian EE cohort

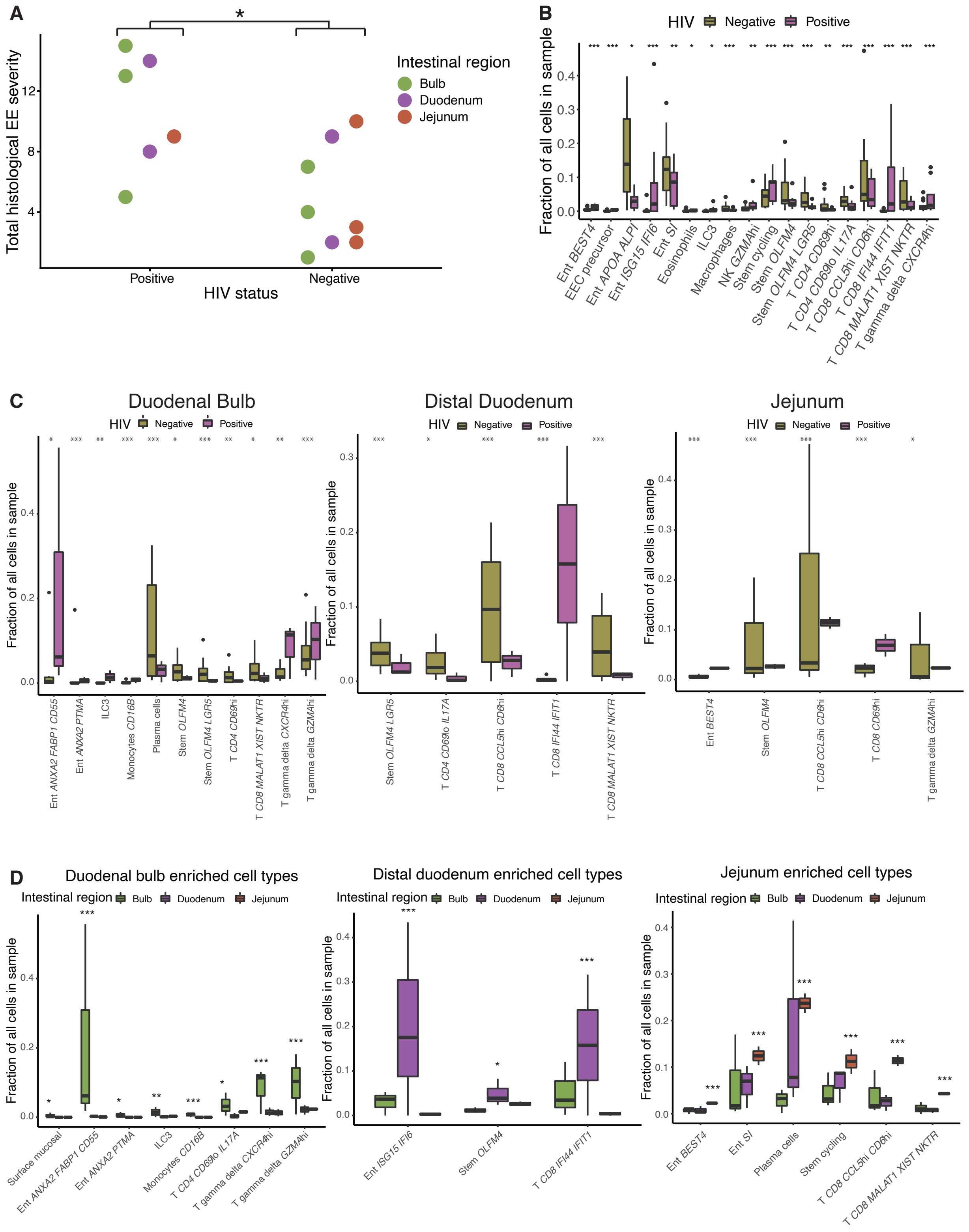

A
Total histological EE severity
HIV status
Positive
Negative
Intestinal region
Bulb
Duodenum
Jejunum
B
HIV Negative Positive
Fraction of all cells in sample
Ent BEST4
EEC precursor
Ent APOA ALPI
Ent ISG15 IFI6
Ent SI
Eosinophils
ILC3
Macrophages
NK GZMAhi
Stem cycling
Stem OLFM4
Stem OLFM4 LGR5
T CD4 CD69hi
T CD4 CD69lo IL17A
T CD8 CCL5hi CD6hi
T CD8 IFI44 IFIT1
T CD8 MALAT1 XIST NKTR
T gamma delta CXCR4hi
C
Duodenal Bulb
Distal Duodenum
Jejunum
HIV Negative Positive
Fraction of all cells in sample
Ent ANXA2 FABP1 CD55
Ent ANXA2 PTMA
ILC3
Monocytes CD16B
Plasma cells
Stem OLFM4
Stem OLFM4 LGR5
T CD4 CD69hi
T CD8 MALAT1 XIST NKTR
T gamma delta CXCR4hi
T gamma delta GZMAhi
Stem OLFM4 LGR5
T CD4 CD69lo IL17A
T CD8 CCL5hi CD6hi
T CD8 IFI44 IFIT1
T CD8 MALAT1 XIST NKTR
Ent BEST4
Stem OLFM4
T CD8 CCL5hi CD6hi
T CD8 CD69hi
T gamma delta GZMAhi
D
Duodenal bulb enriched cell types
Distal duodenum enriched cell types
Jejunum enriched cell types
Intestinal region Bulb Duodenum Jejunum
Fraction of all cells in sample
Surface mucosal
Ent ANXA2 FABP1 CD55
Ent ANXA2 PTMA
ILC3
Monocytes CD16B
T CD4 CD69lo IL17A
T gamma delta CXCR4hi
T gamma delta GZMAhi
Ent ISG15 IFI6
Stem OLFM4
T CD8 IFI44 IFIT1
Ent BEST4
Ent SI
Plasma cells
Stem cycling
T CD8 CCL5hi CD6hi
T CD8 MALAT1 XIST NKTR

**Figure S7: Variation in EE biology associated with HIV infection**

**A,** EE HIV-positive samples had significantly higher total EE histological severity scores than EE HIV-negative samples ($p < 0.05$; Wilcoxon test).

**B,** Cell subsets with a significant change in fractional abundance between HIV-positive and HIV-negative samples across all EE samples (*,adj. $p < 0.05$; **, adj. $p < 0.01$; ***, adj. $p < 0.001$).

**C,** Cell subsets with a significant change in fractional abundance between HIV-positive and HIV-negative samples within EE samples from each intestinal region (*,adj. $p < 0.05$; **, adj. $p < 0.01$; ***, adj. $p < 0.001$).

**D,** Within only HIV-positive EE samples, cell subsets with a significant change increase in fractional abundance in each intestinal region (*,adj. $p < 0.05$; **, adj. $p < 0.01$; ***, adj. $p < 0.001$).

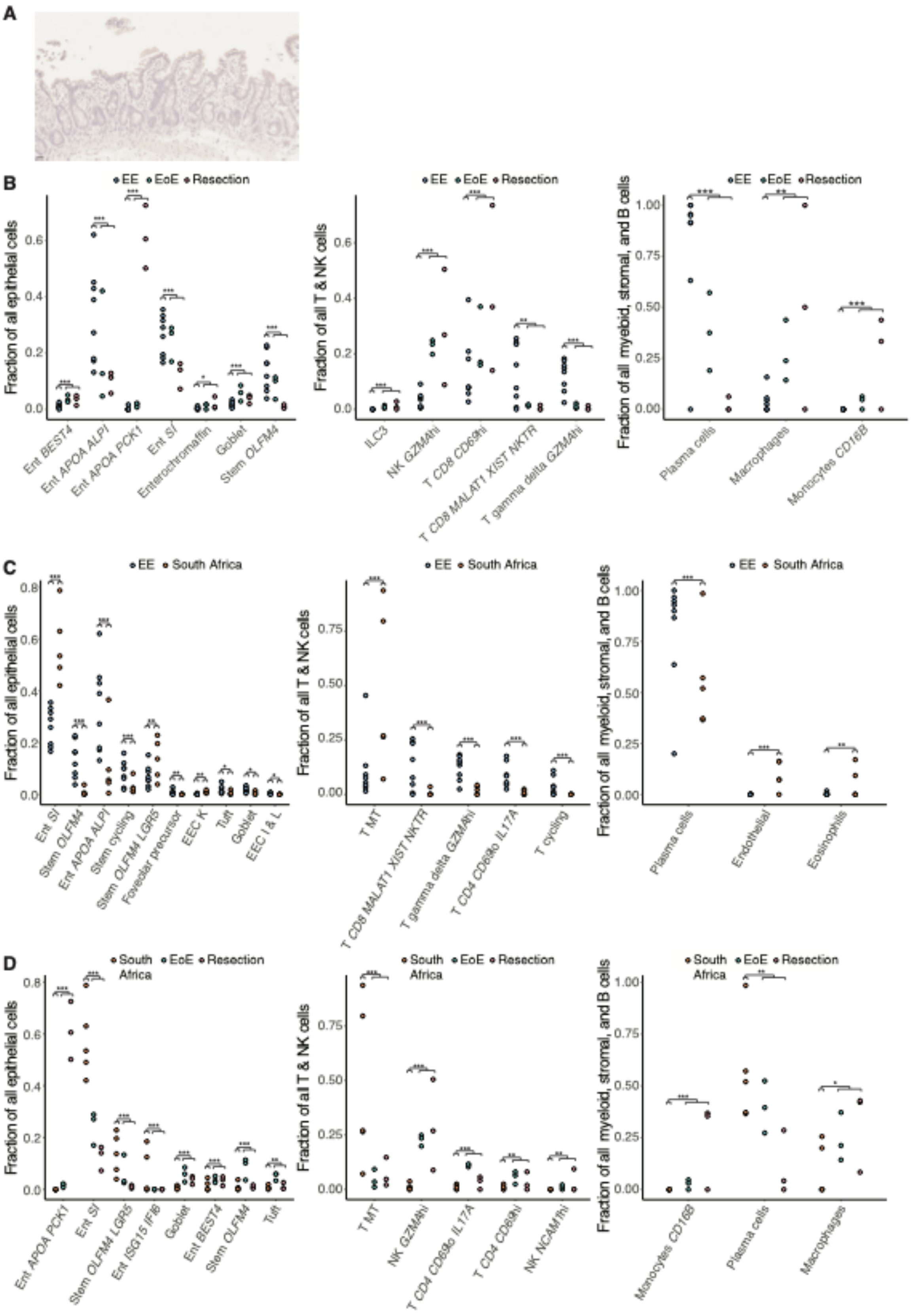
A
B
EE
EoE
Resection
Fraction of all epithelial cells
Ent BEST4
Ent APOA ALPI
Ent APOA PCK1
Ent SI
Enterochromaffin
Goblet
Stem OLFM4
Fraction of all T & NK cells
ILC3
NK GZMAhi
T CD8 CD69hi
T CD8 MALAT1 XIST NKTR
T gamma delta GZMAhi
Fraction of all myeloid, stromal, and B cells
Plasma cells
Macrophages
Monocytes CD16B
C
EE
South Africa
Fraction of all epithelial cells
Ent SI
Stem OLFM4
Ent APOA ALPI
Stem cycling
Stem OLFM4 LGR5
Foveolar precursor
EEC K
Tuft
Goblet
EEC I & L
Fraction of all T & NK cells
T MT
T CD8 MALAT1 XIST NKTR
T gamma delta GZMAhi
T CD4 CD69lo IL17A
T cycling
Fraction of all myeloid, stromal, and B cells
Plasma cells
Endothelial
Eosinophils
D
South Africa
EoE
Resection
Fraction of all epithelial cells
Ent APOA PCK1
Ent SI
Stem OLFM4 LGR5
Ent ISG15 IFI6
Goblet
Ent BEST4
Stem OLFM4
Tuft
Fraction of all T & NK cells
T MT
NK GZMAhi
T CD4 CD69lo IL17A
T CD4 CD69hi
NK NCAM1hi
Fraction of all myeloid, stromal, and B cells
Monocytes CD16B
Plasma cells
Macrophages

**Figure S8: Samples from South African participants display features of EE**

**A,** H&E staining from a biopsy taken from a participant at the same clinical site in South Africa where the participants in this study were profiled.

**B,** Cell subsets with significant shifts in relative abundances between the Zambian EE cohort and both U.S. cohorts (*,adj. $p < 0.05$; **, adj. $p < 0.01$; ***, adj. $p < 0.001$; Fischer's exact test).

**C,** Cell subsets with significant shifts in relative abundances between the Zambian EE cohort and the South African cohort (*,adj. $p < 0.05$; **, adj. $p < 0.01$; ***, adj. $p < 0.001$; Fischer's exact test).

**D,** Cell subsets with significant shifts in relative abundances between the South African cohort and both U.S. cohorts (*,adj. $p < 0.05$; **, adj. $p < 0.01$; ***, adj. $p < 0.001$; Fischer's exact test).

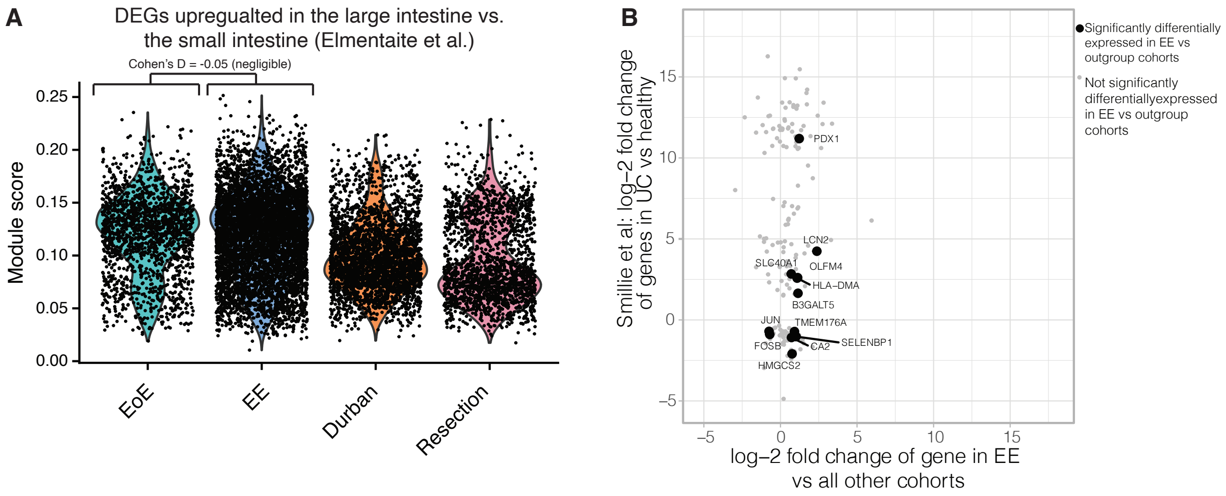
A
DEGs upregualted in the large intestine vs.
the small intestine (Elmentaite et al.)
Cohen's D = -0.05 (negligible)
Module score
0.25
0.20
0.15
0.10
0.05
0.00
EoE
EE
Durban
Resection
B
Smillie et al: log–2 fold change
of genes in UC vs healthy
log–2 fold change of gene in EE
vs all other cohorts
15
10
5
0
–5
–5
0
5
10
15
PDX1
LCN2
SLC40A1
OLFM4
HLA–DMA
B3GALT5
JUN
TMEM176A
FOSB
CA2
SELENBP1
HMGCS2
Significantly differentially expressed in EE vs outgroup cohorts
Not significantly differentiallyexpressed in EE vs outgroup cohorts

**Figure S9: Comparing EE epithelial cells to signatures from existing scRNA-seq datasets of the gastrointestinal tract**

**A,** Violin plots of module scores for a set of genes enriched in the large intestine relative to the small intestine for all HIV-negative distal duodenal samples from each participant cohort.

**B,** Scatterplot of the genes differentially expressed in the EE epithelium relative to all three control cohorts and the genes differentially expressed in the large intestine with ulcerative colitis relative to the healthy large intestine.

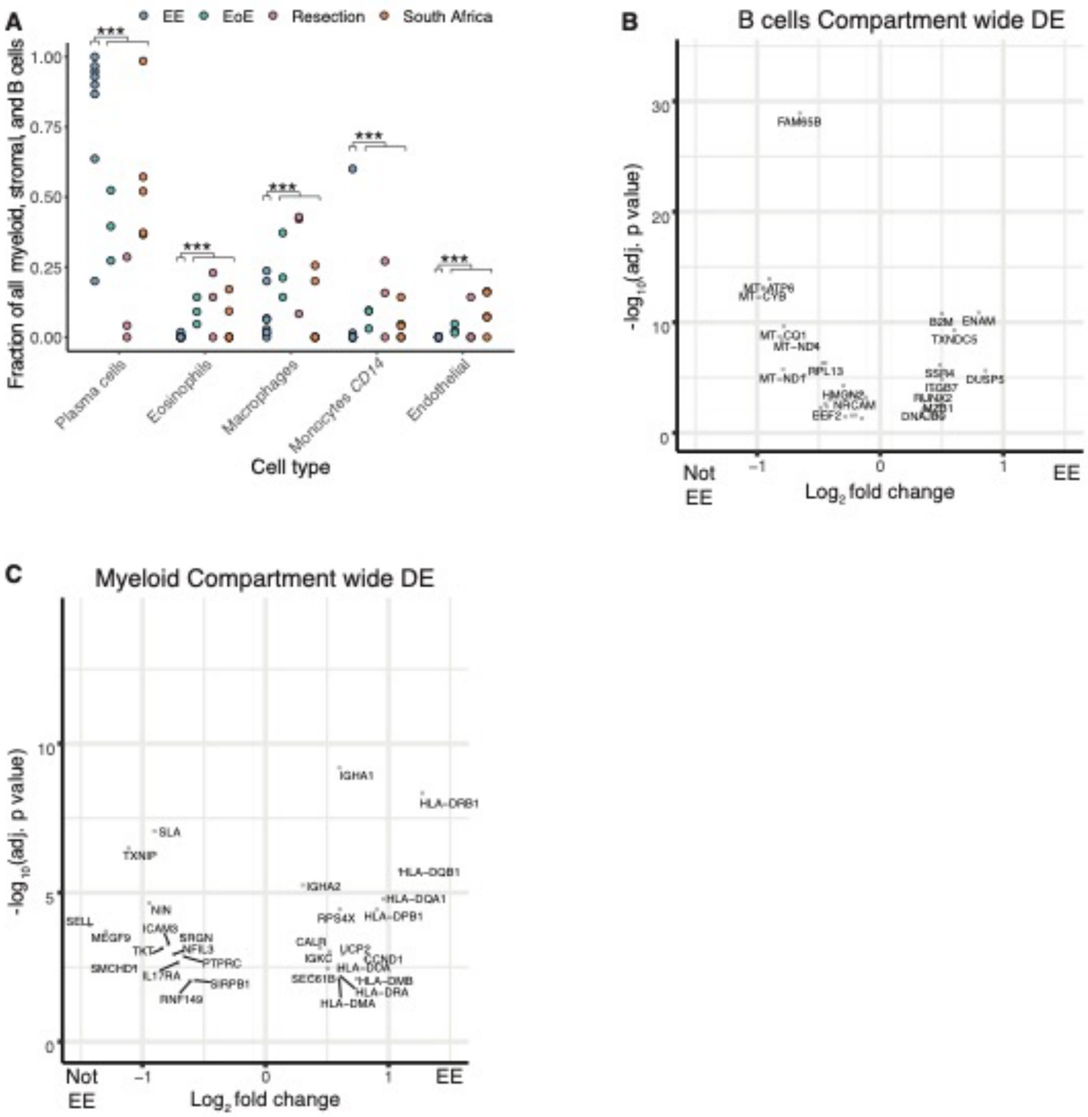
A
EE
EoE
Resection
South Africa
Fraction of all myeloid, stromal, and B cells
1.00
0.75
0.50
0.25
0.00
***
Plasma cells
Eosinophils
Macrophages
Monocytes CD14
Endothelial
Cell type
B
B cells Compartment wide DE
$-\log_{10}$(adj. p value)
30
20
10
0
FAM65B
MT−ATP6
MT−CYB
MT−CO1
MT−ND4
MT−ND1
RPL13
HMGN2
NRCAM
EEF2
B2M
ENAM
TXNDC5
SSR4
DUSP5
ITGB7
RUNX2
MZB1
DNAJB9
−1
0
1
Not EE
EE
$Log_2$ fold change
C
Myeloid Compartment wide DE
$-\log_{10}$(adj. p value)
10
5
0
IGHA1
HLA−DRB1
SLA
TXNIP
IGHA2
HLA−DQB1
HLA−DQA1
NIN
RPS4X
HLA−DPB1
SELL
MEGF9
ICAM3
SRGN
TKT
NFIL3
PTPRC
SMCHD1
IL17RA
SIRPB1
RNF149
CALR
UCP2
IGKC
CCND1
HLA−DOA
SEC61B
HLA−DMB
HLA−DRA
HLA−DMA
−1
0
1
Not EE
EE
$Log_2$ fold change

**Figure S10: Cell subset relative abundance and differential expression changes in B cells and myeloid cells between EE and controls**

**A,** Cell subsets with significant shifts in relative abundances between EE and control cohorts (*,adj. $p < 0.05$; **, adj. $p < 0.01$; ***, adj. $p < 0.001$; Fischer's exact test).

**B,** Genes differentially expressed in all B cells in EE relative to controls.

**C,** Genes differentially expressed in all myeloid cells in EE relative to controls.

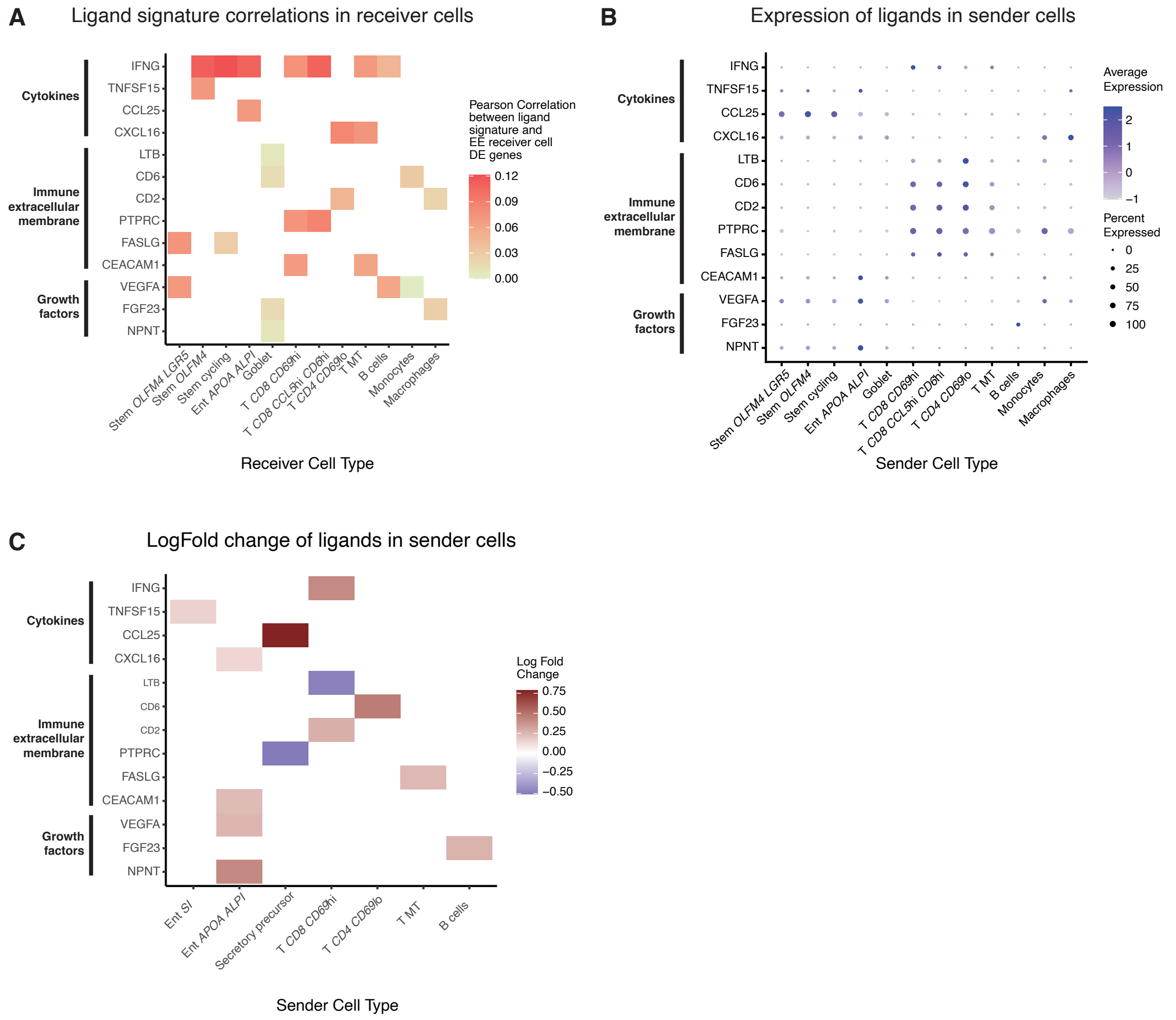
A
Ligand signature correlations in receiver cells
Cytokines
Immune extracellular membrane
Growth factors
IFNG
TNFSF15
CCL25
CXCL16
LTB
CD6
CD2
PTPRC
FASLG
CEACAM1
VEGFA
FGF23
NPNT
Stem OLFM4 LGR5
Stem OLFM4
Stem cycling
Ent APOA ALPI
Goblet
T CD8 CD69hi
T CD8 CCL5hi CD6hi
T CD4 CD69lo
T MT
B cells
Monocytes
Macrophages
Receiver Cell Type
Pearson Correlation between ligand signature and EE receiver cell DE genes
0.12
0.09
0.06
0.03
0.00
B
Expression of ligands in sender cells
Sender Cell Type
Average Expression
2
1
0
−1
Percent Expressed
0
25
50
75
100
C
LogFold change of ligands in sender cells
Ent SI
Ent APOA ALPI
Secretory precursor
T CD8 CD69hi
T CD4 CD69lo
T MT
B cells
Sender Cell Type
Log Fold Change
0.75
0.50
0.25
0.00
−0.25
−0.50

**Figure S11: NicheNet analysis of cell-cell signaling in EE relative to controls**

**A**, Pearson correlation between ligand NicheNet ligand response signatures and the differential gene expression signature in each subset.

**B**, Expression levels of the genes for NicheNet ligands in all cell subsets.

**C**, Log-fold change of the NicheNet ligands in all cell subsets.

A

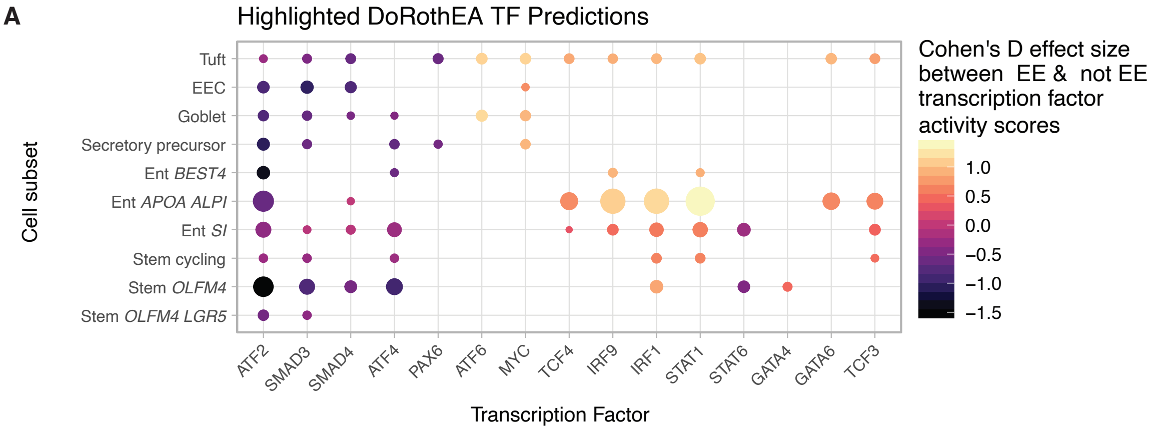


B

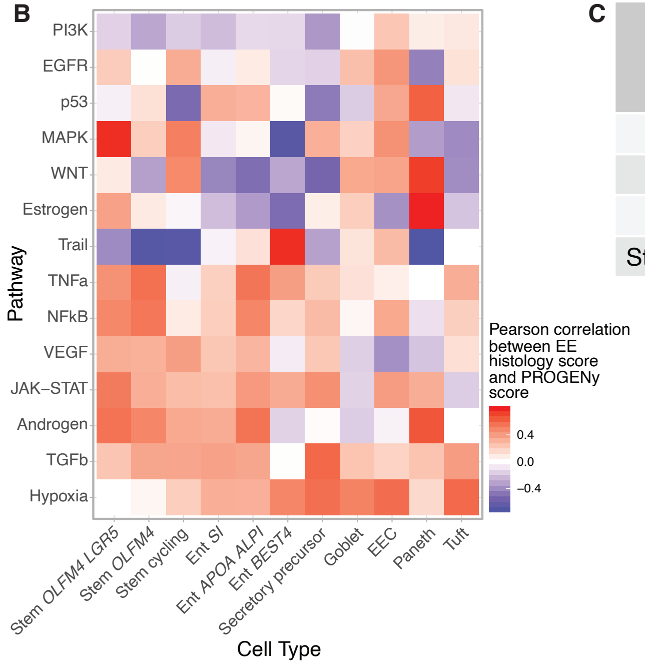


C

| **Cell subset** | **Pearson correlation between histological severity and cell cycle gene module** |
|---|---|
| All stem subsets | 0.09735497 |
| Stem cycling | 0.05718507 |
| Stem *OLFM4* | -0.08540477 |
| Stem *OLFM4 LGR5* | -0.1234793 |

**Figure S12: Additional characterization of epithelial signaling changes in the EE cohort relative to the U.S. cohorts**

**A,** Transcription factors with DoRothEA predicted activity varying between EE and controls (adj. $p < 0.05$).

**B**, Heatmap of Pearson correlations between PROGENY pathway scores and EE histological severity among EE patients.

**C,** Pearson correlation between EE histological severity and proliferation module score within stem cell subsets from EE patients.

#### SUPPLEMENTARY TABLES

**Tables S1-S9 have been uploaded as excel files in the Auxiliary Supplementary Materials section**

### SUPPLEMENTARY MATERIALS AND METHODS

#### Tissue digestion for scRNA-seq

Single-cell suspensions were obtained using a modified version of a previously published protocol*(22)*, described below in detail. Biopsies were rinsed in 30 mL of ice cold HBSS, then transferred to 10 mL of epithelial cell solution (HBSS, 10 mM EDTA, Pen / Strep, 10 mM HEPES, 2% FCS) and incubated for 15 minutes at 37 degrees Celsius at 120 rpm. Samples were transferred to sit on ice for 10 minutes and then vortexed for 10 seconds, after which the tissue was rinsed in 30 mL HBSS before being transferred to1 mL of epithelial digestion mix (2% FCS, 10 mM HEPES. Pen/Strep, 20 ug/ml gentamicin, 100 ug/mL liberase TM, 100 ug/mL DNAse I) in a 1.5 mL Eppendorf tube and spun down at 800g for 2 minutes and resuspended in TrypLE express enzyme [ThermoFisher 12604013] for 5 minutes at 37 degrees Celsius followed by gentle trituration with a P1000 pipette and spun down at 800 g for 2 minutes. Next, after aspiration, the pellet was resuspended in ACK lysis buffer [ThermoFisher A1049201] and put on ice for 3 minutes. Cells were then spun down at 800g for 2 minutes and resuspended in 1 mL of epithelial cell solution and placed on ice for 3 minutes before triturating with a P1000 pipette. Cells were then filtered into a new Eppendorf tube through a 40 uM cell filter [Falcon/VMR 21008-949]. The epithelial cell fraction–which passed through the filter–was spun down at 800g for 2 minutes and then resuspended in 200 mL and put on ice, while the tissue remaining on the filter was added to 5 mL of enzymatic digestion mix at 37 degrees Celsius for 30 minutes at 120 rpm, after which it was quenched with 80 uL of 0.5M EDTA) and placed on ice for 5 minutes. Samples were typically fully dissociated at this step and after gentle trituration with a P1000 pipette filtered through a new 40 uM strainer into a new 50 mL conical tube and rinsed with HBSS to 30 mL total volume, and spun down at 400g for 10 minutes. All but 1 mL was then aspirated off of the 50 mL conical, after which the pellet was resuspended in this 1 mL, which was then transferred to a 1.5 mL Eppendorf. This was spun down at 800g for 2 minutes and then resuspended in 1 mL of ACK lysis butter and put on ice for 3 minutes. Cells were then spun down at 800 g for 2 minutes and resuspended in 200 mL of epithelial cell solution and placed on ice, produced the lamina propria cell fraction. The lamina propria and epithelial fractions were then counted using a hemocytometer.

#### Single-cell RNA-seq with Seq-Well $S^3$ experimental details

We used Seq-Well $S^3$ (Ref. *(20)*), for our single-cell profiling, for which full methods are available on the Shalek Lab website (www.shaleklab.com). Briefly, 16,000 cells (8,000 from the epithelial fraction from tissue digestion and 8,000 from the lamina propria fraction from tissue digestion) were loaded onto a functionalized polydimethylsiloxane (PDMS) array preloaded with ~80,000 uniquely-barcoded mRNA capture beads (Chemgenes; MACOSKO-2011-10). The array was then sealed with a hydroxylated polycarbonate membrane with pore sizes of 10 nm, thereby enabling buffer exchange (and thereby cellular lysis and mRNA transcript hybridization to beads) while retaining biological molecules confined within each well. Beads were then removed from each well, reverse transcribed, treated with Exonuclease I (New England Biolabs; M0293M), and mixed with 0.1 M NaOH for 5 minutes at room temperature to denature the mRNA-cDNA hybrid product on the bead. Second strand synthesis was then performed with a PCR mastermix (40 uL 5x maxima RT buffer, 80 uL 30% PEG8000 solution, 20uL 10 mM DNTPS, 2uL 1 mM dn-SMART oligo, 5 uL Klenow Exo-, and 53 ul of DI ultrapure water) which was incubated with the beads for 1 hour at 37 ºC with end over end rotation. PCR amplification was then preformed using KAPA HiFi PCR Mix (Kapa Biosystems KK2602). After PCR amplification whole transcriptome products were isolated via two rounds of SPRI purification using Ampure Spri beads (Beckman Coulter, Inc.) at both 0.6x and 0.8x volumetric ratio. Sequencing libraries were then generated using the Nextera Tagmentation method on a total of 800 pg of pooled cDNA library per sample. This product was then purified via two rounds of SPRI at 0.6 and 0.8x ratios, producing library size with an average distribution of 500-750 basepairs, as determined using the Aligent hsD1000 Screen Tape System (Aligent Genomics). Arrays were sequenced on an Illumina Next-Seq at the Ragon Institute. The read structure was paired end, with read 1 starting at a custom read 1 primer containing 21 bases with a 12 bp cell barcode and an 8 bp unique molecular identifier (UMI) and read 2 consisting of 50 bases containing transcript information. Sequencing read alignment and demultiplexing was performed on the cumulus platform*(80)* using snapshot 6 of the Drop-seq pipeline previously described in*(52)*, resulting in a cell barcode by UMI digital gene expression (DGE) matrix.

**Data QC and clustering overview**

Prior to clustering, the DGE matrices were filtered to remove cellular barcodes with less than 250 unique genes, with more than 50,000 UMIs, with over 50% of UMIs mapping to mitochondrial genes, and identified as doublets via the Scrublet algorithm*(83)* (which was ran separately for each Seq-Well array). Due to the multiple intestinal regions sampled and multiple patient cohorts in this study, different samples contained different distributions of cell types. In particular, unique subsets of cells were present in duodenal bulb samples, in samples from HIV-positive individuals, and in samples from uninvolved tissue from patients with pancreatic cancer. We tested the Seurat V3 integration approach*(84)* to see if we could correct for batch effects between samples while preserving differences driven by biology (i.e. differences driven by the region of the tissue sampled or the disease state of the patient). Initial testing revealed that integration incorrectly merged biologically distinct cell types in the duodenal bulb with secretory lineage cells from the second part of the duodenum and the jejunum. Furthermore, integration removed differences in clustering driven by HIV status that were seen without integration, indicating that integration may have been over correcting the dataset and minimizing biological differences driven by disease biology. Thus, rather than running an integration method for batch correction, we adopted an existing pipeline for automated iterative clustering of single-cell data (pipeline described in detail in the next methods section) that has been shown to identify batch effects without collapsing distinct rare cell types*(26, 81)*.

**Workflow for iterative clustering**

1. We ran the clustering pipeline on the full filtered DGE matrix and manually identified base clusters with co-expression of known mutually exclusive cellular lineage markers and removed these base clusters that correspond to doublet populations.
2. We ran the clustering pipeline separately on the epithelial, T and NK cell, B cell, myeloid, and stromal cellular compartments, generating a hierarchical tree of clusters for each compartment.
3. Known lineage specific genes*(19, 22, 28)* were used to annotate cell types on the clustering tree. The genes used to annotate each subset are available alongside the genes differentially upregulated in each subset in

Supplementary Table 4. If sub clusters of a cluster did not have lineage defining genes as marker genes, then the cell type designation was made at the parent cluster. In over 95%, cell type designations were made at the first or second level of clustering, and lower tiers of clusters (representing transcriptional differences within a cell type) were not annotated as cell types. In some cases, multiple sub clusters expressed marker genes for one cell type and were all given that cell type designation. For example, within an epithelial cluster composed of secretory lineage cells, two clusters that highly expressed *TFF3* and *CLCA1* were merged together as one final goblet cell type.

4. Finally, in one instance, we performed batch correction by identifying populations of cells with high expression of known lineage markers that formed separate sub clusters in the hierarchical clustering tree due to transcriptional differences between disease states and merging the clusters with common high expression of a lineage marker back together. This correction was only carried out on stem cells highly expressing *OLFM4* and *LGR5*, where the clustering pipeline split a tier 1 cluster of stem cells by disease specific expression changes in the AP-1 signalling pathway at tier 2 before splitting by the co-expressed stem marker genes *LGR5* and *SMOC2* at tier 3, producing two separate *LGR5*$^{hi}$ sub clusters at tier 3 with differing levels of AP-1 signalling activation. These two sub clusters were merged together as the Stem *OLFM4*$^{hi}$ *LGR5*$^{hi}$ cell type. Furthermore, although *APOA1* and *APOA4* are markers for mature enterocytes near the villus tips*(85)*, we chose to not merge a cluster of enterocytes predominantly from pancreatic cancer resection samples that exhibited high expression of *APOA1* and *APOA4* with a different cluster of enterocytes expressing *APOA1* and *APOA4* as enterocyte sub types lie on a transcriptional gradient and these two clusters displayed distinctly different expression of *ALPI*–another marker for mature villus enterocytes*(85)*– which raised the possibility that these enterocyte clusters were biologically distinct. We thus did not merge these cell types.

**Iterative clustering pipeline**

Starting with our filtered DGE matrix, cells were iteratively clustered with the following unbiased pipeline built using the Seurat R pacakge*(86)* and adopted from*(26, 81)*. For each starting cluster, normalization and variable gene selection were conducted with SCtransform*(87)*, then dimensional reduction was carried out by running PCA. For starting clusters with less than 500 cells, the number of PCs was chosen with the JackStraw function in Seurat, while for runtime considerations, for starting clusters with more than 500 cells, the number of PCs was chosen as the elbow on the PCA variance explained plot. Cellular neighbours were calculated using a k.param of 0.5*(number of cells)^0.5. Next clustering was carried out at 40 resolutions evenly spaced between 0.2 to 0.8, and the average silhouette width across cells was calculated for each resolution. After calculating a histogram of the average silhouette scores with (num resolutions)/1.2 bins, the resolution in the top bin of silhouette scores with the smallest average silhouette score in that bin was chosen as the final silhouette score, thereby creating a bias towards under clustering the data. Next, marker genes were calculated for each sub cluster generated with the final resolution using the Wilcoxon test with pre-test thresholds that each gene must have an average log fold change of at least 0.025 between clusters and be expressed in at least 0.2 percent of cells in the cluster that it is a marker for. For each sub cluster, this clustering pipeline was iteratively repeated until sub clusters were reached that did not have at least 10 marker genes with at least an adjusted p value of 0.01 and an average log fold change of 0.2. Altogether, this pipeline generated a hierarchical tree of clusters which then used in conjunction with known lineage markers to annotate cell types as described above.

**Details on module scores for reduced villus height and reduced plasma LPS associated genes**

Gene signatures for reduced villus height and decreased and plasma LPS signatures in Chama et al.*(7)* were obtained by aggregating all genes positively differentially expressed in these conditions in this study. Module scores were generated with the AddModuleScore function in Seurat with default parameters. Cell subsets were then scored for the modules, a Wilcoxon test was used to assess significance, and effect sizes were calculated with Cohen's D.

**Identifying cell subsets varying with histological EE severity across all HIV-negative patients in the Zambian cohort**

Cell subsets significantly associated with histological severity from the Zambian cohort were identified by running Dirichlet Regression, which allows for testing for differences in cell subsets along a continuous dependent variable, with the R package DirichletReg. Across all HIV-negative samples, relative abundances were regressed against the total EE histological severity score and small intestinal region, and an adjusted p value was generated for the association between EE severity and the relative abundance of each subset. Other cell subsets co-varying with the significantly associated subsets from Dirichlet regression were visualized by calculating the Pearson correlation between all subset relative abundances across patients and then hierarchically clustering the resulting correlations with Ward's method using the ComplexHeatmap R package. To identify cell subsets significantly associated with surface mucosal cells, sample labels were permuted 10,000 times and the Pearson correlations of each subset with surface mucosal cells were recalculated to form null distributions for the association between each cell type and surface mucosal cells. Original Pearson correlation values within the top 5% of values of the respective null distribution for each cell subset were designated as significant correlations.

**Further details for analyses identifying biological features that distinguish HIV-negative EE from U.S. control cohorts**

Compositional testing: Cell subset differences were calculated using the leave-one-out approach described to identify changes in HIV status and intestinal region.

Differential expression: Differentially expressed genes for each cell subset and for the epithelial and immune compartments as a whole were found with a Wilcoxon test implemented in the Seurat FindAllMarkers function with the minimum log-fold change threshold set to 0.1 and the minimum percent of cells within a subset expressing a gene set to 0.025.

PROGENy: PROGENy scores for signalling pathway activities were calculated using the progeny R package using 500 genes to generate the model matrix and all other parameters set to default as suggested by the tutorial vignette for applying PROGENy to scRNA-seq data.

DoRothEA: DoRothEA scores for upstream transcription factor activities were calculated using the dorothea R package following the parameters suggested in the Bioconductor vignette for applying DoRothEA to scRNA-seq data.

Module scoring: To identify changes in proliferative capacity in stem cells in EE relative to stem cells in the U.S. control cohorts, we examined a list of genes previously identified as upregulated in cycling human cells*(52)*. To score for a signature of tissue residency in T and NK cells, we used a previously identified signature*(88)* for tissue resident memory T cells consisting of the genes *CD69, ITGAE, ITGA1, IL2, IL10, CXCR6, CXCL13, KCNK5, RGS1, CRTAM, DUSP6, PDCD1, and IL23R.* T cell signatures for activation were obtained from a previous scRNA-seq atlas of T cell activation*(67)*. For all gene signatures, module scores were generated with the AddModuleScore function in Seurat with default parameters. Cell subsets were then scored for the modules, a Wilcoxon test was used to assess significance, and effect sizes were calculated with Cohen's D.

Comparison with large intestine gene signature from Elmentaite et. al.: Intestinal scRNA-seq data from adults was downloaded from Elmentaite et al and the Find Markers function with default parameters in Seurat was used to find genes upregulated in the large intestine vs the small intestine.

Comparison with Ulcerative colitis gene signature from Smillie et al: A gene signature comparing the inflamed colon with ulcerative colitis was obtained from the supplemental information in Smillie et al.

**Predicting ligand-receptor interactions with NicheNet**

NicheNet signatures for changes in ligand-receptor interactions between EE and the control cohorts were generated from the differentially expressed genes generated as described above. All other parameters in NicheNet were kept the same as in the NicheNet tutorial "Perform NicheNet analysis starting from a Seurat object".

**Metagenomic mapping of H. pylori reads with Kraken2**

Bam files from sequencing were classified using Kraken2 with the following Terra pipeline from the Broad institute: https://dockstore.org/workflows/github.com/broadinstitute/viral-pipelines/classify_kraken2:master?tab=files. The following inputs were used. Kraken2_db_tgz: "gs://pathogen-public-dbs/v1/kraken2-broad-20200505.tar.zst". Krona_taxonomy_db_tgz: "gs://pathogen-public-dbs/v1/krona.taxonomy-20200505.tab.zst". The outputted results for the samples were then aggregated across all bacterial species using the merge_metagenomics pipeline on the Terra platform from the Broad institute: https://viral-pipelines.readthedocs.io/en/latest/merge_metagenomics.htmls
